# Inhibition of mitochondrial respiration impairs nutrient consumption and metabolite transport in human retinal pigment epithelium

**DOI:** 10.1101/2020.05.11.086827

**Authors:** Rui Zhang, Abbi L Engel, Yekai Wang, Bo Li, Weiyong Shen, Mark C Gillies, Jennifer Chao, Jianhai Du

**Affiliations:** Department of Ophthalmology, West Virginia University, Morgantown, WV 26506; Department of Biochemistry, West Virginia University, Morgantown, WV 26506; Save Sight Institute, Sydney Medical School, University of Sydney, Sydney, NSW 2000, Australia; Department of Ophthalmology, University of Washington, Seattle, WA 98109

**Keywords:** mitochondrial respiration, metabolism, retinal pigment epithelium, metabolites, glucose, amino acids, nucleotides, ketone bodies

## Abstract

Mitochondrial respiration in mammalian cells not only generates ATP to meet their own energy needs but also couples with biosynthetic pathways to produce metabolites that can be exported to support neighboring cells. However, how defects in mitochondrial respiration influence these biosynthetic and exporting pathways remains poorly understood. Mitochondrial dysfunction in retinal pigment epithelium (RPE) cells is an emerging contributor to the death of their neighboring photoreceptors in degenerative retinal diseases including age-related macular degeneration. In this study, we used targeted-metabolomics and ^13^C tracing to investigate how inhibition of mitochondrial respiration influences the intracellular and extracellular metabolome. We found inhibition of mitochondrial respiration strikingly influenced both the intracellular and extracellular metabolome in primary RPE cells. Intriguingly, the extracellular metabolic changes sensitively reflected the intracellular changes. These changes included substantially enhanced glucose consumption and lactate production, reduced release of pyruvate, citrate and ketone bodies, and massive accumulation of multiple amino acids and nucleosides. In conclusion, these findings reveal a metabolic signature of nutrient consumption and release in mitochondrial dysfunction in RPE cells. Testing medium metabolites provides a sensitive and noninvasive method to assess mitochondrial function in nutrient utilization and transport.

---

The retinal pigment epithelium (RPE) maintains the health of the neural retina by performing several critical functions, including nutrient transport, phagocytosis of the outer segments and secretion of cytokines (41). These RPE-specific processes rely on active energy metabolism. Recent evidence indicates that dysfunctional RPE mitochondrial bioenergetics is a crucial factor for many retinal degenerative diseases, including age-related macular degeneration (AMD), one of the leading causes of blindness (9, 21, 24). Inhibition of mitochondrial metabolism specifically in RPE is sufficient to induce AMD-like retinal degeneration in mice (19, 30, 50). RPE cells from AMD donors have impaired mitochondrial metabolism (20, 23). However, the detailed mechanism of how impaired mitochondrial metabolism in RPE results in photoreceptor degeneration remains elusive.

Mitochondrial respiration is the central process in energy metabolism. Reducing equivalents NADH and FADH_2_, generated by the tricarboxylic acid (TCA) cycle, donate electrons to complex I to II in the electron transport chain (ETC), and oxygen is finally consumed to produce ATP through ATP synthase (complex V). Besides being a primary generator of energy, mitochondrial respiration through the TCA cycle also provide critical biosynthetic pathways to generate glucose, fatty acid, ketone bodies, and non-essential amino acids (22, 36). Growing evidence shows that RPE mitochondria are crucial for the metabolic communication between RPE and outer retina by synthesizing nutrients to support retinal health (45, 47, 48). Each day, approximately 10% of photoreceptor outer segments are shed and phagocytosed by RPE (33). RPE mitochondria are capable of converting the phagocytosed lipids into ketone bodies and recycling them back to fuel photoreceptors (1, 39). Furthermore, the retina generates a massive amount of lactate by the Warburg effect (13, 27). A recent study indicated that RPE mitochondria may use lactate released from photoreceptors as an energy source to maintain their function, thus preserving the majority of glucose for photoreceptors (28). Moreover, we recently showed that RPE mitochondria prefer to metabolize nutrients such as proline into intermediates that are exported towards the retinal side (10). Nevertheless, how mitochondrial dysfunction impacts RPE nutrient utilization and secretion of metabolites towards retina is not well defined.

Mitochondrial function is typically assessed by measuring oxygen consumption in isolated mitochondria or live cells by sequentially adding substrates and inhibitors of mitochondrial complexes. The major drawbacks of these techniques are: (1) cells cannot be re-used after the assay, and (2) the activities of mitochondrial biosynthetic pathways are not measured. Primary human fetal or adult RPE cultures are excellent disease models since they maintain many properties similar to native RPE. These primary cultures are expensive and labor-intensive, so a noninvasive method to quantify mitochondrial function for these cells is highly desirable.

In the present study, we used targeted-metabolomics approaches to investigate how inhibition of mitochondrial complexes influences the intracellular and extracellular metabolome in primary cultured human RPE cells. Our findings demonstrate that inhibition of mitochondrial complexes causes early and unique changes in medium metabolites, RPE mitochondrial function is critical for nutrient synthesis, and quantification of metabolites in the media may provide a novel method for measuring mitochondrial metabolism.

## EXPERIMENTAL PROCEDURES

### Reagents

Mitochondrial respiration inhibitors, culture media, and other reagents are listed in detail in **Table S5**.

### Primary RPE cell culture

Human RPE culture was generated from human donors without known ocular diseases as previously described (16). The protocols for this culture and sample preparation were approved by the Institutional Review Board from both the University of Washington and the West Virginia University. RPE was initially plated in a 12-well or 35mm tissue culture plates coated with Growth Factor Reduced (GFR) Matrigel® Matrix (Corning). The cells were cultured in Minimum Essential Medium Alpha (MEMα) medium, supplemented with 5% (vol/vol) fetal bovine serum (FBS), N1 Medium Supplement, hydrocortisone, triiodo-thyronine, taurine, nonessential amino acids and a penicillin-streptomycin solution (See details in **Table S5**). 10uM of Y-27632 dihydrochloride was included for the first 1-2 weeks of culturing. At confluency, the media was changed to a 1% (vol/vol) FBS supplemented media without Y-27632 dihydrochloride. RPE cells were passaged to a 12-well plate at a density of 400,000-550,000 cells/well for experiments. The RPE cells were cultured for at least 4 to attain maturity before use in experiments.

### Cell Treatment and sample preparation

The mature RPE cells were treated with 1μM piericidin A, 1 μg/ml antimycin A, or 5μM oligomycin in regular 1% supplemented RPE media described above. An equivalent volume of diluent DMSO was added as vehicle control (1μL/ml). 50 µL of media was collected at 1h, 6h and 24h. Cells were harvested at 24h by quickly rinsing with a cold 0.9% NaCl solution, and adding 100μL of 80% methanol pre-cooled at −20°C. Cells were scraped on dry ice and rinsed with an additional 100μL of 80% methanol. The cells were transferred to a microcentrifuge homogenized and centrifuged to extract metabolites as describe (14).

For labeling with [^13^C_6_]-glucose, piericidin, antimycin, and oligomycin or control DMSO were added at the same concentrations to clear DMEM media (Gibco A144301) without, then supplemented with 1% FBS and penicillin-streptomycin, and 5mM [^13^C_6_]-glucose. RPE cells were changed into the DMEM media with DMSO or inhibitors. The media and cells were collected in the same way as mentioned above.

### Metabolite analysis with liquid chromatography-mass spectrometry (LC MS) and gas chromatography-mass spectrometry

Metabolite preparation and analysis with LC MS and GC MS were performed as previously reported (14, 48). Medium samples were centrifuged to remove debris and 10 µL of supernatant was mixed with 40 µl of cold methanol to extract metabolites. Supernatants containing aqueous metabolites from both media and cells were dried at 4°C and analyzed by LC MS or GC MS. LC MS used a Shimadzu LC Nexera X2 UHPLC coupled with a QTRAP 5500 LC MS/MS (AB Sciex). An ACQUITY UPLC BEH Amide analytic column (2.1 × 50 mm, 1.7 μm, Waters) was used for chromatographic separation. Each metabolite was tuned with standards for optimal transitions. The extracted MRM peaks were integrated using MultiQuant 3.0.2 software (AB Sciex). Table S6 lists the detailed parameters for the measured metabolites. For GC MS, the samples were derivatized by methoxyamine and N-tertbutyldimethylsilyl-N-methyltrifluoroacetamide, and analyzed by the Agilent 7890B/5977B GC MS system with a DB-5MS column (30 m × 0.25 mm × 0.25 μm film). Mass spectra were collected from 80–600 m/z under selective ion monitoring mode. The detailed parameters were listed in **Table S6**. The data was analyzed by Agilent MassHunter Quantitative Analysis Software and natural abundance was corrected by ISOCOR software.

### Statistical Analysis

Multivariate analysis and the number of changed metabolites were analyzed with PCA and One-way ANOVA respectively, using MetaboAnalyst 4.0 (https://www.metaboanalyst.ca/). Univariate statistical analyses were performed by Graphpad Prism 8.0 (GraphPad Software, Inc. La Jolla, CA). Data are presented as mean±standard error. Values of p<0.05 were considered significantly different.

## RESULTS

### Inhibition of mitochondrial respiration severely disrupts nutrient consumption and utilization in RPE cells

To study how mitochondrial dysfunction affects metabolites, we used piericidin, antimycin and oligomycin to inhibit mitochondrial complex I, complex III and complex V respectively in fully differentiated primary RPE cells, and then collected culture media at 1h, 6h and 24h and cells at 24h to analyze metabolites by mass spectrometry (**Fig 1A**). To evaluate the toxicity of these inhibitors, we analyzed lactate dehydrogenase (LDH) activity in the media since dead cells release LDH into the culture media. We did not observe changes in LDH activity at 1h, 6h, and 24h for all inhibitors except antimycin, which increased LDH activity only at 24h (**Fig S1**). To study whether the overall metabolites in the media were different between groups treated with control (dimethyl sulfoxide, DMSO) or inhibitors, we performed a multivariate analysis of the metabolomics data with principal component analysis (PCA). The scores plot showed that the metabolites with three inhibitors in the media at 1h overlapped with those from the control, but the metabolites with inhibitors were well separated from the control in media and cells at 6h and 24h after treatment (**Fig 1B-E**). Although the oligomycin group had a distinctive profile in the cells, it was not separate from the groups with the other two inhibitors in the media. Among the 126 metabolites we detected, 11∼43 extracellular metabolites were significantly changed from 1h to 24h, and 118 intracellular metabolites were significantly altered (**Fig 1F, Table S1-4**). Pathway analysis showed that the changed extracellular metabolites were enriched in amino acid metabolism, glycolysis and TCA cycle, nucleotide metabolism, ketone bodies and acylcarnitines, NAD metabolism and urea cycle (**Fig 1G**). Mitochondrial complexes are critical for the production of ATP, which could be stored as phosphocreatine (PCr). As expected, high-energy metabolites, including ATP, ADP, and PCr were significantly attenuated by these inhibitors, confirming that mitochondrial energy metabolism was severely impaired in the primary RPE cells (**Fig 1H**).

**Figure 1.**
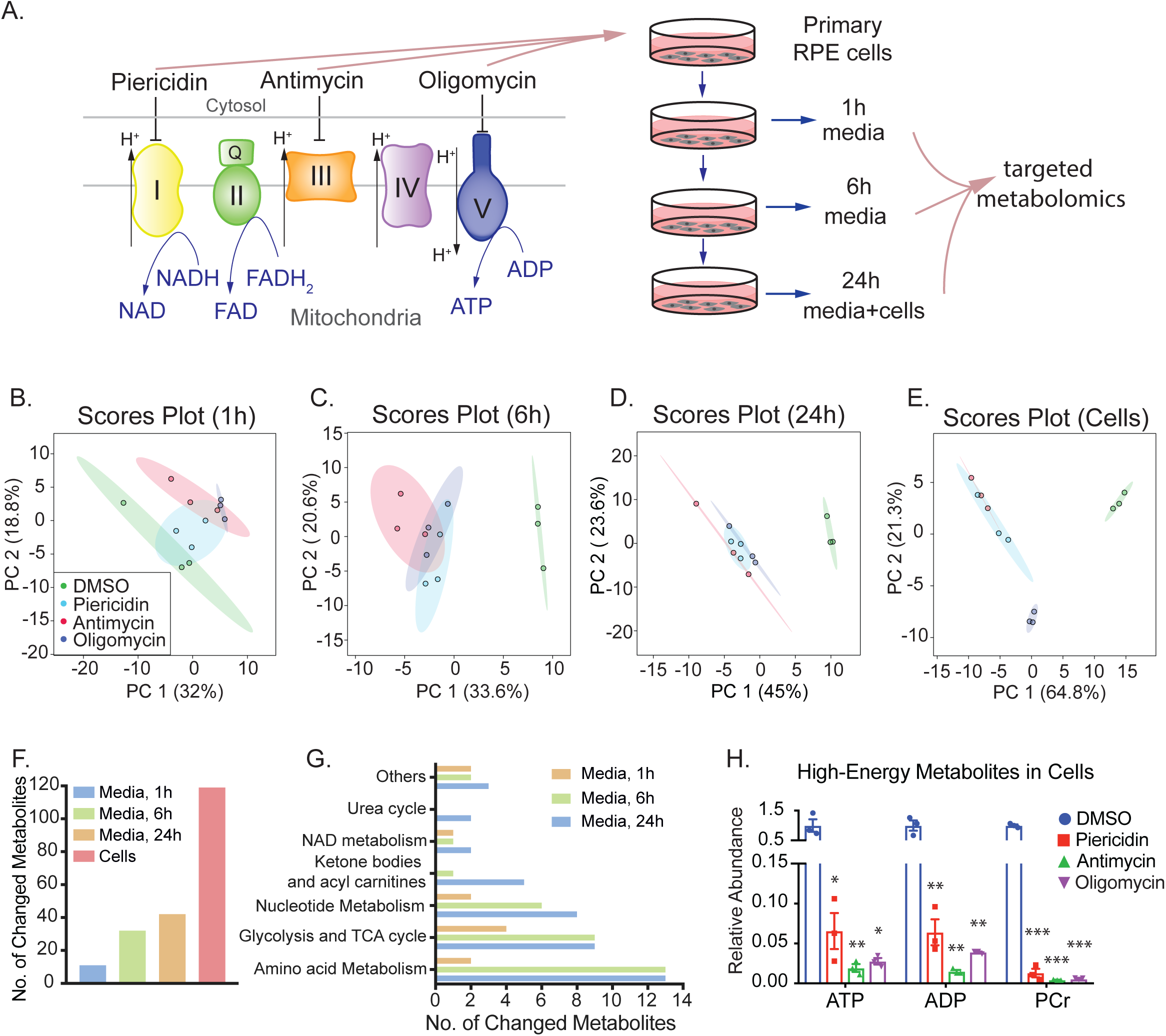
Inhibition of mitochondrial metabolism changes both intracellular and extracellular metabolites in human RPE cells. (**A)** Experimental design for quantifying intracellular and extracellular metabolites caused by mitochondrial dysfunction. Mitochondrial electron transport chain shuttles electrons from NADH and FADH to O_2_ to produce ATP. We inhibited Complex I, III or V with their specific inhibitors, piericidin, antimycin or oligomycin, respectively, in primary cultured RPE cells to investigate the impact of mitochondrial dysfunction on intracellular metabolites. Media and cells were collected at different time points as indicated and quantified by targeted metabolomics. **(B-E)** Scores plots of metabolites in cells and medium at 1h, 6h, and 24h by principal component analysis. (**F)** The number of changed metabolites in the medium at 1h, 6h, 24h, and in RPE cells. **(G)** The number of changed metabolites in different pathways in the medium at 1h, 6h, 24h. (**H)** Mitochondrial inhibitors decreased the levels of high-energy metabolites in cells. N=3. *P<0.05, **P<0.01, *P<0.001 vs. the cells treated with DMSO. PCr, phosphocreatine.

### Inhibition of mitochondrial respiration disrupts glucose utilization and metabolism

Glucose is an important nutrient to produce energy and intermediates through glycolysis and the TCA cycle (**Fig 2A**). After 24h, mitochondrial inhibitors almost completely depleted intracellular glucose but significantly elevated intracellular lactate. However, most intracellular glycolytic intermediates upstream of lactate were decreased, especially by piericidin and antimycin (**Fig 2B**). Intracellular TCA cycle intermediates including citrate, isocitrate and α-KG were decreased but oxaloacetate (OAA) accumulated, indicating that citrate synthesis is inhibited (**Fig 2C**). Antimycin and oligomycin, but not piericidin, substantially increased intracellular succinate, confirming that the inhibition of complexes is specific because succinate does not need complex I for its oxidation (**Fig 2C**). Similar to the intracellular changes, glucose, pyruvate, citrate and isocitrate dropped 2-10 fold as early as 6h in the media while lactate and OAA were increased by all inhibitors (**Fig 2D-H, 2M**). Similarly, extracellular succinate was increased by antimycin and oligomycin at 24h (**Fig 2J**). These results support the concept that medium metabolites are important indicators for the metabolic status inside cells. However, the extracellular levels of α-KG, fumarate and malate had different patterns of change (**Fig 2I, 2K-L**), indicating the mechanisms that maintain metabolites across the cell membranes are more complex for these metabolites.

**Figure 2.**
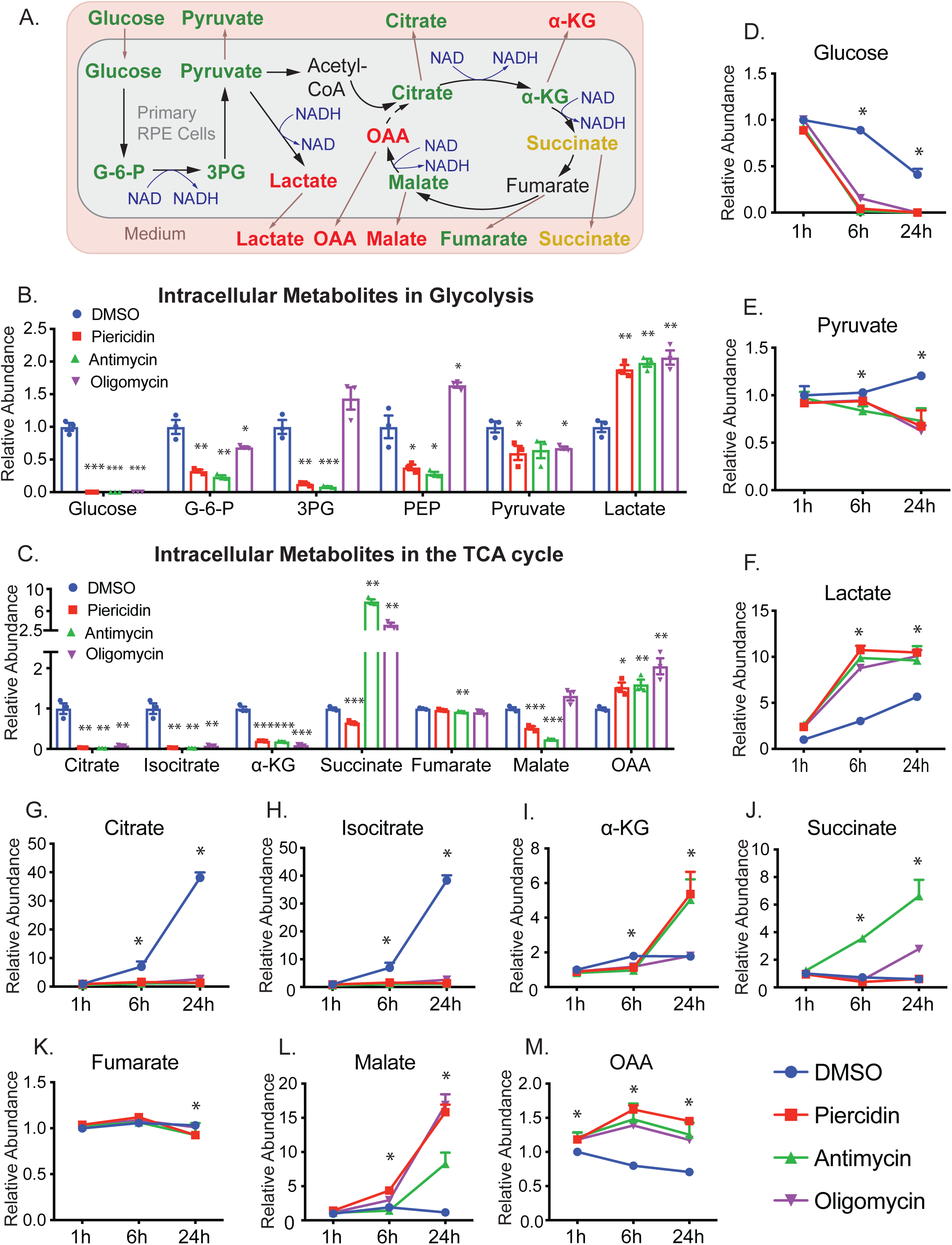
Inhibition of mitochondrial metabolism impairs intracellular and extracellular metabolites in glycolysis and the TCA cycle. (**A)** A schematic of the impact of mitochondrial dysfunction on glucose metabolism. Green represents a decrease, red represents an increase, and orange represents mixed changes by different inhibitors. (**B-C)** Significantly changed intracellular metabolites in glycolysis and the TCA cycle in the RPE cells. N=3. *P<0.05, **P<0.01, *P<0.001 vs. the cells treated with DMSO. (**D-M)** The changed extracellular metabolites in glucose metabolism. N=3. *P<0.05 vs. DMSO. G-6-P, glucose-6-phosphate; 3PG, 3-phosphoglycerate; PEP, phosphoenolpyruvate.

### Inhibition of mitochondrial respiration impairs the utilization of ^13^C glucose

The intermediates in glycolysis and the TCA cycle can be derived from multiple nutrients. To ask whether the changes come from glucose, we used [^13^C_6_]-glucose to trace glucose metabolism and analyzed ^13^C-labeled metabolites with GC-MS (**Fig 3A**). To avoid the interference of supplements such as glucose and pyruvate, we switched the media to DMEM without glucose. After the addition of 5 mM [^13^C_6_]-glucose in DMEM, more than 90% of intracellular pyruvate and lactate, and 25-80% of intracellular TCA intermediates were ^13^C-labeled (**Fig S2**). Except for lactate, the enrichment (percentage of labeling) of all the intermediates was reduced when mitochondrial metabolism was inhibited (**Fig S2**). Similarly, the levels of ^13^C-labeled intracellular pyruvate and mitochondrial intermediates were substantially diminished by mitochondrial inhibitors, while lactate was slightly increased or unchanged (**Fig 3B-C**). These results indicate that the inhibition of mitochondrial complexes may block the oxidation of NADH and activate the conversion of pyruvate into lactate. As expected, the ratio of total lactate to pyruvate, an indicator of cytosolic NADH, was increased 5-10 fold by these mitochondrial inhibitors (**Fig S3**). Consistently, the exported ^13^C-labeled lactate in the media increased as early as 1h and had increased further at 6h and 24h, but ^13^C pyruvate release decreased (**Fig 3D-E**), suggesting that the medium lactate and pyruvate are sensitive indicators of mitochondrial metabolism. We found RPE cells had high efflux of glucose-derived citrate and α-KG, but the inhibition of mitochondrial respiration entirely blocked this efflux (**Fig 3F-G**), consistent with the intracellular changes. However, intracellular and extracellular succinate, fumarate and malate changed in different patterns after mitochondrial inhibition (**Fig 3H-J**), indicating other factors or mechanisms are involved in regulating their release.

**Figure 3.**
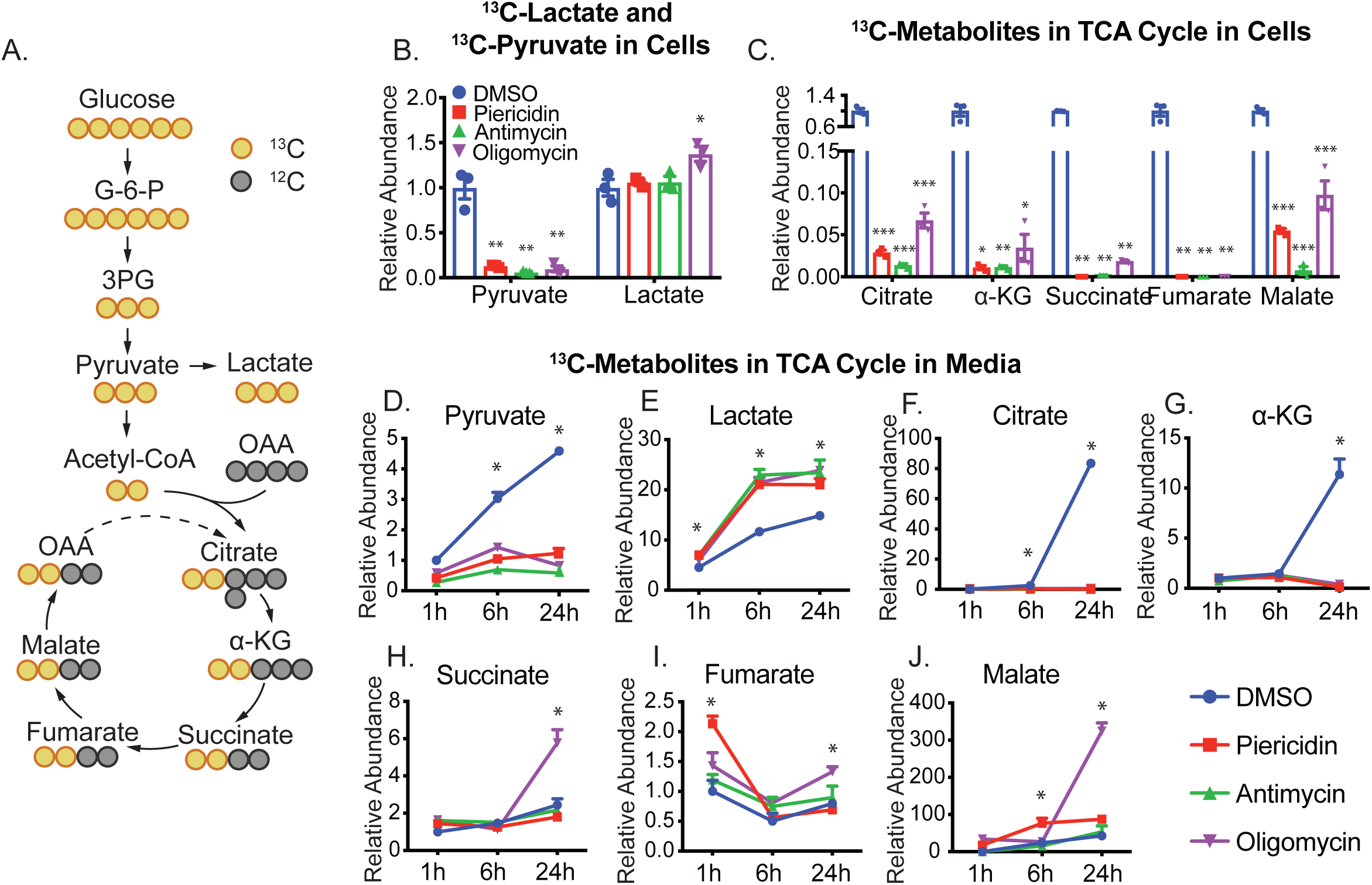
Inhibition of mitochondrial metabolism impairs glucose utilization. (**A)** A schematic for ^13^C labeling in intermediates in glycolysis and TCA cycle from [^13^C_6_]-glucose (yellow circles). OAA, oxaloacetate. **(B-C)** The relative abundance of ^13^C-labelled intracellular metabolites over the group treated with DMSO. **(D-J)** The relative abundance of ^13^C-labelled extracellular metabolites over the group treated with DMSO. N=3. *P<0.05, **P<0.01, ***P<0.001 vs. the groups treated with DMSO.

### Inhibition of mitochondrial respiration impairs the amino acid metabolism

The regular RPE culture medium is typically supplemented with abundant amino acids. The inhibition of mitochondrial respiration caused widespread changes of 44 intracellular amino acids and 13 extracellular amino acids (**Fig 4, Fig S3-4, Table S1-4**). Choline and proline were the two most sensitive and dramatically changed amino acids. With mitochondrial inhibition, both amino acids in the media were significantly increased at 1h and had accumulated 5-20 fold by 24h (**Fig 4C-D**). Consistently with changes in the medium, intracellular choline and proline also accumulated 5-12 fold (**Fig 4A**). These results suggest that choline and proline are sensitive amino acid markers for RPE mitochondrial metabolism. We found healthy RPE cells time-dependently consumed branch-chain amino acids (BCAAs) including leucine, isoleucine and valine from the media. However, both intracellular and extracellular BCAAs were elevated with the inhibition of mitochondrial metabolism (**Fig 4A, E-G**). These data suggest mitochondrial dysfunction blocks the utilization of BCAAs.

**Figure 4.**
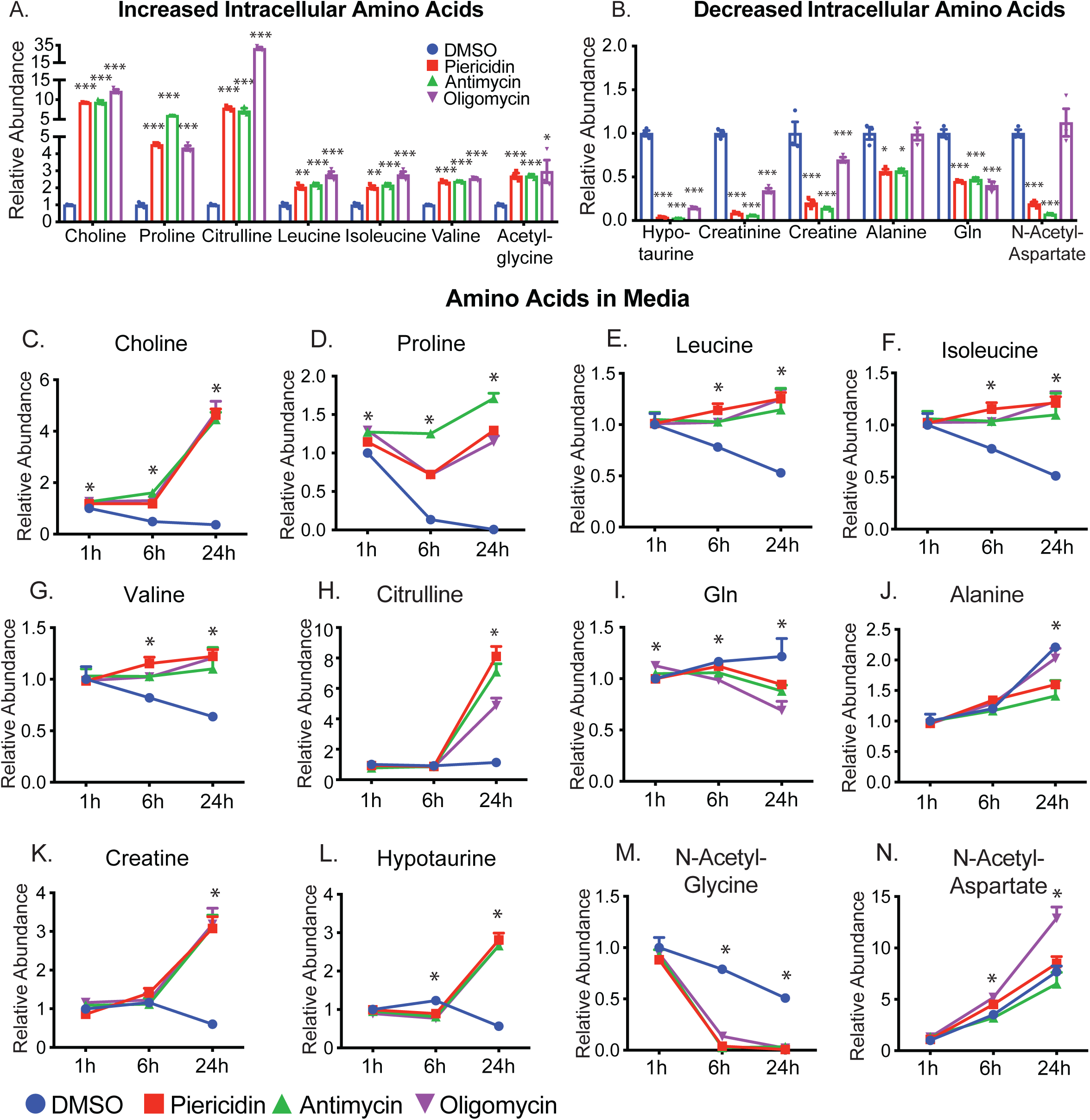
Inhibition of mitochondrial metabolism impairs amino acid metabolism and the urea cycle. **(A-B)** The relative abundance of significantly increased or decreased intracellular amino acids. **(C-N)** The relative abundance of amino acids in the media at different time points. N=3. *P<0.05, **P<0.01, ***P<0.001 vs. the groups treated with DMSO. Gln, glutamine.

Although the urea cycle is restricted to the liver, several enzymes in the urea cycle are ubiquitous in mammalian cells to regulate aspartate, citrulline and arginine metabolism (**Fig S4**). Both intracellular and extracellular citrulline accumulated substantially (**Fig 4A, 4H**). Ornithine and aspartate (**Fig S4**) were increased or unchanged, but arginosuccinate was decreased by both piericidin and antimycin (**Fig S4**). These results suggest arginosuccinate synthase, the enzyme that catalyzes the conversion of citrulline into arginosuccinate, might be inhibited by the lack of ATP.

Glutamine and alanine decreased in both cells and media by mitochondrial inhibitors, suggesting that their consumption is enhanced, or their release is inhibited (**Fig 4B, 4I-J**). Changes of most other amino acids in media were opposite or inconsistent with their changes in cells such as creatine, hypotaurine, N-acetyl-glycine, and N-acetyl-aspartate (**Fig 4A-B, Fig 4K-N, Fig S3-S4**). These disparities suggest that efflux or influx of these amino acids are affected differently by the inhibition of mitochondrial respiration.

### Inhibition of mitochondrial respiration impairs the metabolism of ketone bodies and acylcarnitines

RPE has active lipid metabolism to produce and release ketone bodies such as 3-hydroxybutyrate (3-HB) (1, 39) (**Fig 5A**). Consistently, RPE cells time-dependently released 3-HB into the media. However, all three inhibitors almost completely shut down the production and release of 3-HB (**Fig 5A-C**), indicating that fatty acid oxidation is inhibited. Fatty acids use acylcarnitine shuttle into mitochondria for β-oxidation through forming acyl-CoAs. The intracellular levels of acetyl-carnitine (C2), propionyl-carnitine (C3), butyryl-carnitine (C4), hexanoyl-carnitine (C6) and myristoyl-carnitine (C14) were significantly decreased, supporting that inhibition of mitochondrial respiration resulted in impaired fatty oxidation (**Fig 5D**). Propionyl-carnitine, isobutyryl-carnitine, and methylbutyroyl-carnitine could be produced by the oxidation of BCAAs. The reduction of these acyl-carnitines confirms our findings in **Fig 4** that the oxidation of BCAAs is impaired by mitochondrial inhibition. Paradoxically, extracellular acyl-carnitines remained unchanged or accumulated by mitochondrial inhibitors, especially, oligomycin (**Fig 5E-G**). In sum, these data suggest that mitochondrial dysfunction inhibits fatty acid oxidation and ketone body production.

**Figure 5.**
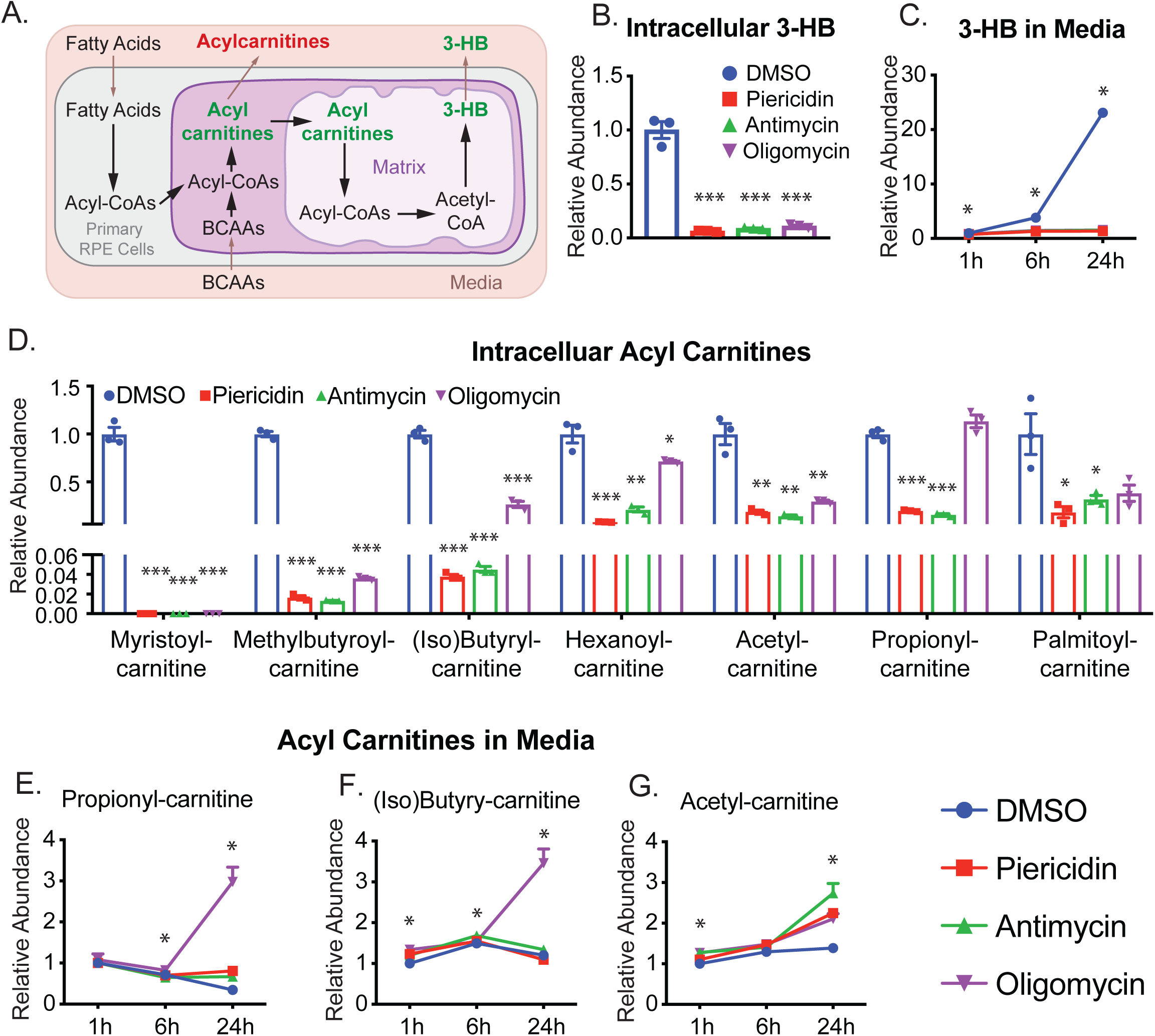
Inhibition of mitochondrial metabolism impairs the metabolism of ketone body production and acylcarnitines. (**A**) A schematic for the metabolism of ketone body and acylcarnitines. (**B-C**) Both intracellular and extracellular 3-hydroxybutyrate (3-HB) was significantly reduced after the inhibition of mitochondrial metabolism. (**D**) Inhibition of mitochondrial metabolism decreased the levels of intracellular acylcarnitines. (E-H) The relative abundance of extracellular acylcarnitines. N=3. *P<0.05, **P<0.01, ***P<0.001 vs. the groups treated with DMSO.

### Inhibition of mitochondrial respiration impairs purine and pyrimidine metabolism

Purine and pyrimidine could synthesize from *de novo* or salvage pathways using glucose and amino acids as precursors or catabolized nucleosides as precursors (**Fig 6A-B**). The inhibition of mitochondrial respiration massively accumulated metabolites from purine degradation such as guanine, guanosine, hypoxanthine and xanthine (**Fig 6C, 6E-J, Fig S5**). These purine metabolites increased by 5-20 fold in both cells and media, whereas some upstream purine intermediates such as ribulose-5-phosphate (R-5-P), inosine monophosphate (IMP) and adenosine monophosphate (AMP) were substantially diminished (**Fig 6A, Fig 6C-J**). These results strongly suggest that inhibition of mitochondrial respiration accelerates the catabolism of purine metabolites and blocks purine synthesis. In a similar pattern, we found metabolites in pyrimidine catabolism including 3-aminoisobutyric acid (BAIBA), cytidine, cytosine, uracil and β-alanine were accumulated in cells or media, but UDP was severely depleted (**Fig 6B-D, Fig 6K-N, Fig S5)**. Impressively, BAIB accumulation in the media started at 1h and reached more than five folds over control at 24h. (**Fig 6M)**. Cytidine was not detected in the control media at 1h but significantly increased in the media with inhibitors at 6h and 24h (**Fig 6K**). Taken together, we found inhibition of mitochondrial metabolism enhances the catabolism of purine and pyrimidine. Nucleotides such as hypoxanthine and BAIBA may serve as sensitive and robust medium indicators for mitochondrial dysfunction.

**Figure 6.**
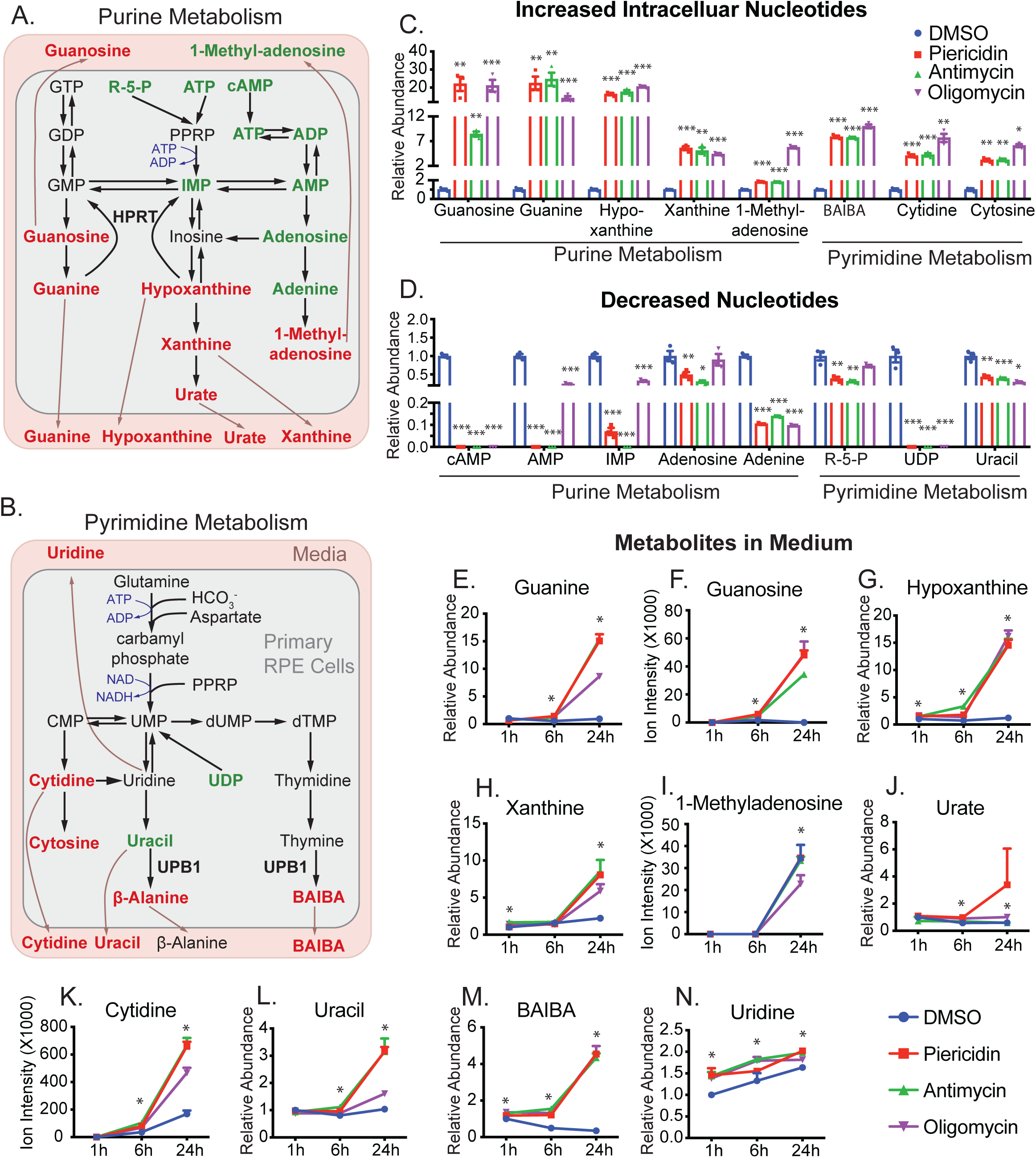
Inhibition of mitochondrial metabolism impairs nucleotide metabolism. (**A-B**) Schematics for purine and pyrimidine metabolism. The metabolites are colored to represent relative abundance by the inhibition of mitochondrial metabolism (red for the increase, green for decrease, and black for no change or not detected). (**C-D**) The relative abundance of significantly changed nucleotides by the inhibition of mitochondrial metabolism. (E-M) The relative abundance of significantly changed medium metabolites in nucleotide metabolism. N=3. *P<0.05, **P<0.01, *P<0.001 vs. the groups treated with DMSO.

### Inhibition of mitochondrial respiration impairs NAD metabolism

NAD is essential in mitochondrial energy metabolism by transferring protons from glycolysis, TCA cycle and fatty acid oxidation. NAD could be synthesized from tryptophan or nicotinamide (**Fig 7A**). Upon the inhibition of mitochondrial respiration, intracellular NAD and its derivatives, NADH and NADP(H), were depleted. (**Fig 7A-B**). Because NAD(H) and NADP(H) are intracellularly produced and impermeable to cell membranes, they could not be detected in the media. Their precursors, tryptophan and nicotinamide, were significantly decreased in cells (**Fig 7B**). Medium tryptophan was not changed, while medium nicotinamide was increased ∼5 fold at 24h (**Fig 7C**). These findings demonstrate that nicotinamide consumption is blocked by mitochondrial inhibitors. The methylation of nicotinamide into 1-methylnicotinamide (MNA) for excretion is an important degradation pathway to regulate nicotinamide homeostasis (**Fig 7A**). We found healthy RPE cells released a large amount of MNA, ∼20-fold increase in 24h. Interestingly, extracellular MNA was significantly deceased, starting from 1h, consistent with intracellular changes except for oligomycin which increased intracellular NMA (**Fig 7B, 7D**). These results indicate that NAD metabolism is impaired by mitochondrial inhibition, and nicotinamide and MNA are useful media markers for NAD metabolism.

**Figure 7.**
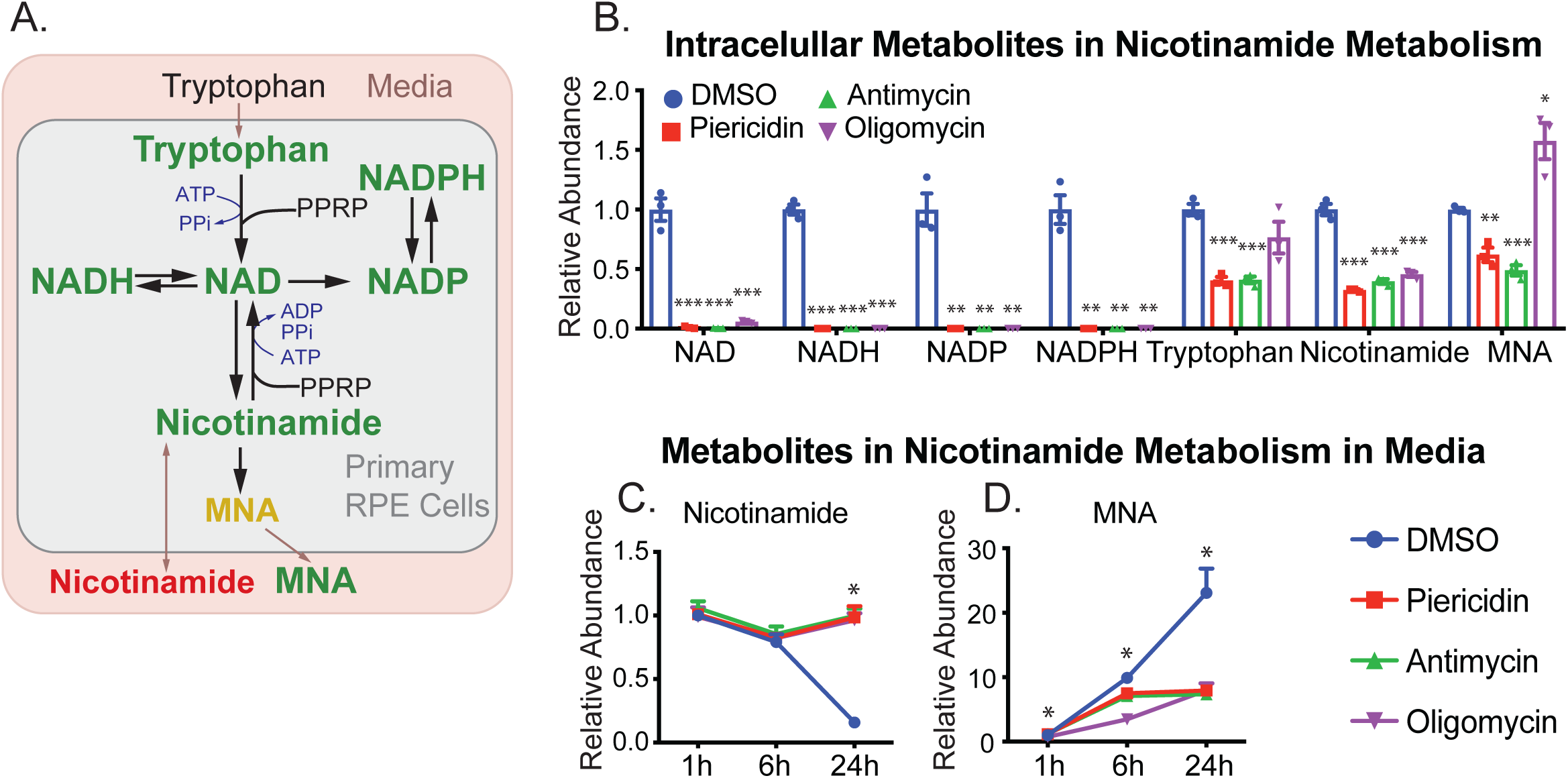
Inhibition of mitochondrial metabolism impairs NAD metabolism. **(A)** A schematic for the impaired NAD metabolism with mitochondria inhibitors. MNA, 1-methylnicotinamide. **(B)** The relative abundance of intracellular NAD intermediates over the groups with DMSO. **(C-D)** The relative abundance of extracellular metabolites in NAD metabolism. N=3. *P<0.05, **P<0.01, ***P<0.001 vs. the groups treated with DMSO.

## DISCUSSION

In this study, we found signature changes of metabolites in the media caused by the inhibition of mitochondrial respiration. These changes include: 1) enhanced glucose consumption and lactate production, 2) reduced release of pyruvate and citrate, 3) accumulated choline, proline and BCAAs, 4) decreased release of 3-HB, 5) accumulated nucleosides from nucleotide degradation, and 6) elevated nicotinamide and MNA (**Fig 8**). Our findings demonstrate that mitochondrial dysfunction disrupts nutrient consumption and secretion, strongly supporting the model that RPE mitochondrial metabolism may actively synthesize nutrients to support the outer retinal metabolism (**Fig 8**).

**Figure 8.**
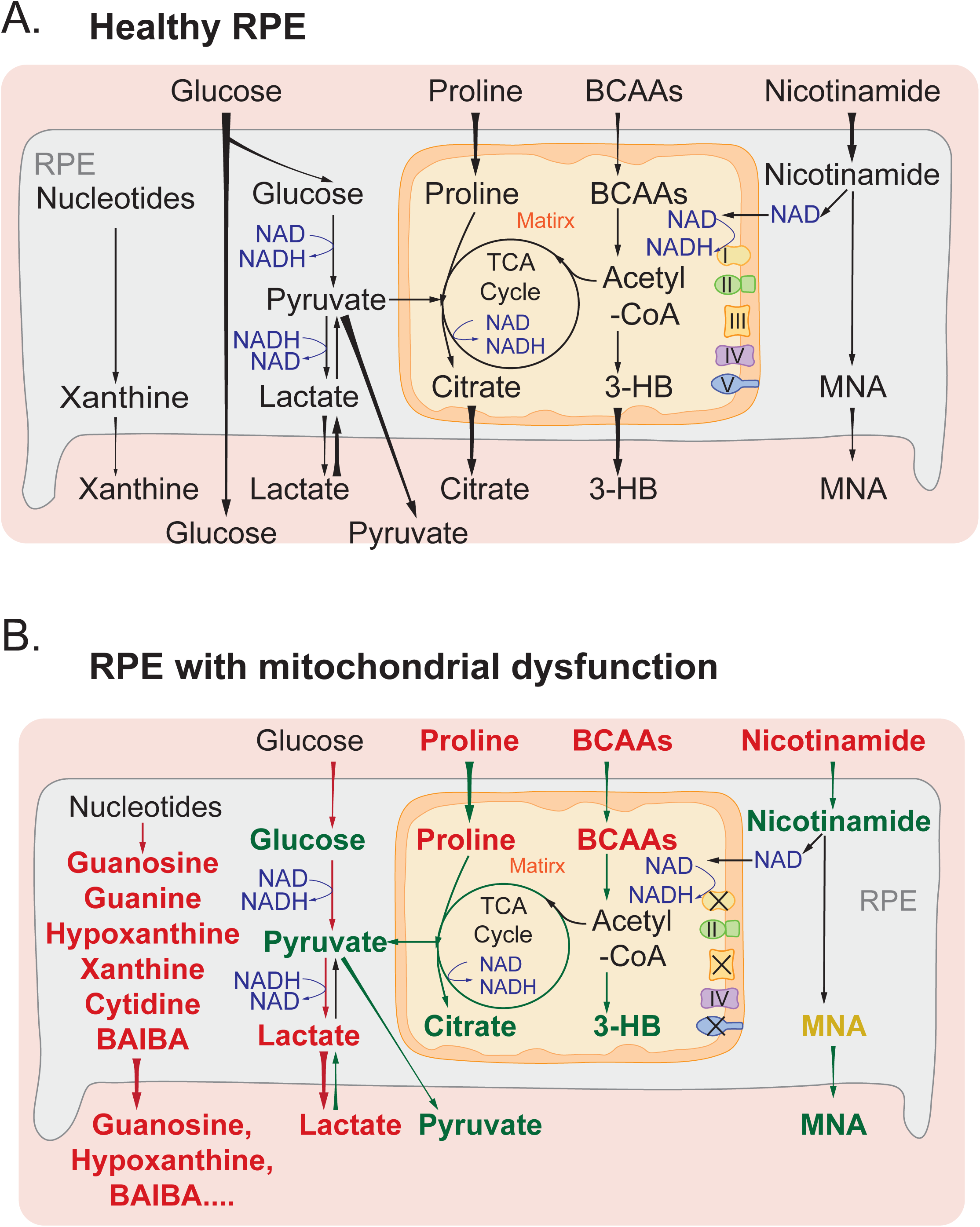
A schematic for the role of mitochondrial metabolism in nutrient consumption and metabolite export. (A) Healthy RPE mitochondria use different nutrients such as glucose, amino acids, lipids and nicotinamide and export mitochondria-derived metabolites including citrate, isocitrate and 3-HB to the outer retina. (B) When mitochondrial respiration is inhibited, RPE cells consume more glucose into lactate but use fewer other fuels, leading to a substantial reduction in exporting glucose, citrate and 3-HB. Moreover, large amounts of lactate and nucleosides such as guanine, guanosine, hypoxanthine, xanthine and BAIBA were exported out of RPE cells.

Why does the inhibition of mitochondrial respiration cause such a broad spectrum of metabolic changes? Mitochondrial respiration accepts electrons from NADH and FADH_2_ to drive the pump of hydrogen to generate ATP and H_2_O by consuming oxygen. The inhibition of mitochondrial respiration by blocking complexes could substantially reduce ATP supply to cause bioenergetics deficit and accumulation of reducing equivalents including NADH/FADH_2_ (11, 42, 44). ATP deficiency would slow or halt many metabolic reactions that require ATP to overcome energy barriers or serve as a cofactor. For example, as ATP is required for the reactions by creatine kinase, arginosuccinate synthase and choline kinase, inhibition of mitochondrial respiration led to depletion of phosphocreatine and accumulation of citrulline and choline (**Fig 1, Fig 4**),. NADH needs to be recycled into NAD to sustain the reactions in glycolysis, TCA cycle and fatty acid oxidation. The ETC is one of the major electron acceptors to regenerate NAD. Both pharmacological inhibitors and genetic deletions of mitochondrial complexes are known to increase NADH/NAD (6, 11, 42, 44). To regenerate NAD from NADH, the cells may use pyruvate to generate lactate through LDH and inhibit NAD-dependent dehydrogenases. Consistently, we have found a substantial increase in lactate and OAA, but a decrease of pyruvate, citrate, isocitrate. The decreased citrate production and enhanced lactate formation could stimulate glycolysis to increase glucose consumption in the RPE cells. Recent studies in other cells also show that inhibition of mitochondrial complex I is sufficient to increase glucose uptake (40, 49). Photoreceptors could uptake citrate and pyruvate to support their mitochondrial metabolism, and export lactate to facilitate glucose transport through RPE (10, 25, 28, 48). The reduced release of citrate and pyruvate from RPE, and accumulation of lactate may interfere with photoreceptor metabolism and disrupt the metabolic ecosystem.

The TCA cycle enzyme succinate dehydrogenase, which is also mitochondrial complex II, accepts electrons from FADH_2_. Succinate decreased from ^13^C glucose (**Fig 2-3**), confirming that mitochondrial oxidation of pyruvate is inhibited. The accumulated succinate in unlabeled RPE culture should come from other sources such as amino acids, fatty acids and pyrimidine. Similar to NAD, FAD needs to be regenerated from FADH_2_ for FAD-dependent dehydrogenases such as acyl-CoA dehydrogenases in fatty acid oxidation and BCAA catabolism (4, 35). Consistently, our findings showed inhibition of mitochondrial respiration reduced acylcarnitines but accumulated BCAAs (**Fig 5**).

Besides being recycled from NADH, NAD needs to be replenished by biosynthesis because multiple pathways consume it such as poly (ADP-ribose) polymerases (PARPs) and sirtuins (3, 5, 34). Activated PARPs can consume large amounts of NAD by transferring ADP-ribose to target proteins (2). We reported previously in primary RPE cells that oxidative stress could deplete cellular NAD, and the inhibition of PARP partially restores NAD to protect oxidative damage (16). Mitochondrial dysfunction could activate PARPs (31), contributing to the depletion of NAD and its derivatives. Furthermore, either *de novo* synthesis from tryptophan or salvage synthesis from nicotinamide requires multiple ATP molecules (34) (**Fig 7A**). The ATP deficiency from inhibition of mitochondrial metabolism could worsen the depletion of cellular NAD. Consistently, nicotinamide was decreased in cells but accumulated in the media (**Fig 7B-C**), confirming that NAD biosynthesis is inhibited. Intriguingly, RPE cells exported large amounts of MNA (**Fig 7D**), methylated nicotinamide by nicotinamide N-methyltransferase (NNMT). Recent studies show NNMT is a novel regulator of energy homeostasis in adipose tissue (17, 29). We reported that MNA is one of several metabolites that are commonly increased in aged ocular tissues (46). The sensitive changes of MNA in the media in our study suggest that medium MNA might be a promising indicator of NAD metabolism and mitochondrial metabolism.

We have reported recently that differentiated mature RPE cells prefer to consume proline as a nutrient and export proline-derived intermediates to be used by the retina (10, 48). In agreement with these findings, healthy RPE cells almost completely consumed proline within 24h. However, complex inhibitors block proline consumption, especially the complex III inhibitor that further elevated proline in the media. To be oxidized, proline is first converted into pyrroline-5-carboxylate (P5C) through a FAD-dependent proline dehydrogenase, and P5C is further oxidized into glutamate by a NAD-dependent P5C dehydrogenase to enter the TCA cycle (37). The ETC inhibitors, especially the inhibitor of complex III that accepts electrons from both FADH_2_ and NADH, could accumulate reducing equivalents and decrease the synthesis of NAD as aforementioned, resulting in the accumulation of proline in the media. As only differentiated RPE that relies on mitochondrial metabolism could utilize proline (48), the medium proline level could be a sensitive marker for RPE mitochondrial metabolism.

One of the striking changes in this study is the massive accumulation of nucleosides in the media from nucleotide metabolism including guanine, guanosine, hypoxanthine, BAIBA, and cytidine. Nucleotide metabolism is finely regulated through synthesis and degradation to maintain DNA integrity and RNA production (32). Nucleosides are also the basic building blocks for high-energy metabolites including ATP, GTP, CTP and UTP. The energy deficit caused by the inhibition of mitochondrial respiration may accelerate the degradation of ADP into AMP, which can activate AMP deaminase to degrade AMP into IMP and hypoxanthine (49). Xanthine oxidoreductase can further oxidize hypoxanthine into xanthine and urate. In this study, AMP and IMP were depleted while hypoxanthine, xanthine, urate, guanine and guanosine were substantially elevated. The salvage pathway through hypoxanthine-guanine phosphoribosyltransferase (HPRT) using hypoxanthine, guanine and xanthine as substrates is the primary pathway for purine synthesis (18) (**Fig 6A**). However, HPRT requires 5-phosphoribosyl 1-pyrophosphate (PRPP) to transfer the 5-phosphoribosyl group. Inhibition of mitochondrial respiration depleted ATP and ribose-5-phosphate, which might decrease PRPP and thus block salvage synthesis. This will ultimately cause further depletion of AMP and IMP. Therefore, our results suggest that inhibition of mitochondrial metabolism increased purine degradation and inhibited its synthesis. Consistently, we have reported that nutrient deprivation causes energy deficient and time-dependent accumulation of purine metabolites including hypoxanthine and xanthine in mouse RPE and retina tissues (46). In photoreceptors, purine metabolism is tightly regulated by light to control the cellular cyclic GMP (cGMP) level (15, 38). Impaired purine metabolism causes inherited retinal degeneration (8, 12). The excessive excretion of purine metabolites from RPE with mitochondrial dysfunction might disturb photoreceptor metabolism and result in retinal degeneration.

We have found a similar accumulation of metabolites in pyrimidine degradation, especially BAIBA. β-ureidopropionase catalyzes the last step in the pyrimidine degradation pathway, producing BAIBA and β-alanine from thymine and uracil, respectively (**Fig 6B**). Both intracellular BAIBA and β-alanine accumulated in the RPE, but in the media, only BAIBA substantially accumulated in a time-dependent manner (**Fig 6M, Fig S5)**. This might be attributed to different pathways in their catabolism. BAIBA is catabolized in the mitochondria into propionyl-CoA, which can be converted into succinyl-CoA to be oxidized through the TCA cycle (43). The accumulation of succinate by the inhibition of mitochondrial respiration should block the degradation of BAIBA to cause its elevation. However, β-alanine can be metabolized into multiple metabolites including acetate, malonate, pantothenic acid, and carnosine (7, 26).

In conclusion, we demonstrate that inhibition of mitochondrial respiration strikingly influences both the intracellular and extracellular metabolome of primary RPE cells, and the extracellular metabolic changes consistently reflect intracellular changes. These findings provide important information on the contribution of mitochondrial dysfunction to the pathogenesis of AMD and other mitochondria-related retinal diseases.

## Acknowledgments

This work was supported by NIH Grants EY026030 (to J.R.C. and J.D.), the Retina Research Foundation (to J.D.), and an unrestricted Grant from Research to Prevent Blindness (J.R.C.).

## Author Contributions

Conceptualization, J.D; Investigation, R.Z., A.L.E., Y.W., J.C., B.L. and J.D.; Writing R.Z., A.L.E., Y.W., W.S., M.C.G., J.C., and J.D; Funding Acquisition J.R.C. and J.D; Supervision, W.S., M.C.G., J.R.C., and J.D.

## REFERENCES

1. Adijanto J, Du J, Moffat C, Seifert EL, Hurle JB, and Philp NJ. The retinal pigment epithelium utilizes fatty acids for ketogenesis. The Journal of biological chemistry 289: 20570–20582, 2014.

2. Alano CC, Garnier P, Ying W, Higashi Y, Kauppinen TM, and Swanson RA. NAD+ depletion is necessary and sufficient for poly(ADP-ribose) polymerase-1-mediated neuronal death. The Journal of neuroscience : the official journal of the Society for Neuroscience 30: 2967–2978, 2010.

3. Ame JC, Spenlehauer C, and de Murcia G. The PARP superfamily. BioEssays : news and reviews in molecular, cellular and developmental biology 26: 882–893, 2004.

4. Barile M, Giancaspero TA, Leone P, Galluccio M, and Indiveri C. Riboflavin transport and metabolism in humans. Journal of inherited metabolic disease 39: 545–557, 2016.

5. Belenky P, Bogan KL, and Brenner C. NAD+ metabolism in health and disease. Trends in biochemical sciences 32: 12–19, 2007.

6. Birsoy K, Wang T, Chen WW, Freinkman E, Abu-Remaileh M, and Sabatini DM. An Essential Role of the Mitochondrial Electron Transport Chain in Cell Proliferation Is to Enable Aspartate Synthesis. Cell 162: 540–551, 2015.

7. Blancquaert L, Baba SP, Kwiatkowski S, Stautemas J, Stegen S, Barbaresi S, Chung W, Boakye AA, Hoetker JD, Bhatnagar A, Delanghe J, Vanheel B, Veiga-da-Cunha M, Derave W, and Everaert I. Carnosine and anserine homeostasis in skeletal muscle and heart is controlled by beta-alanine transamination. The Journal of physiology 594: 4849–4863, 2016.

8. Bowne SJ, Sullivan LS, Blanton SH, Cepko CL, Blackshaw S, Birch DG, Hughbanks-Wheaton D, Heckenlively JR, and Daiger SP. Mutations in the inosine monophosphate dehydrogenase 1 gene (IMPDH1) cause the RP10 form of autosomal dominant retinitis pigmentosa. Human molecular genetics 11: 559–568, 2002.

9. Brown EE, Lewin AS, and Ash JD. Mitochondria: Potential Targets for Protection in Age-Related Macular Degeneration. Advances in experimental medicine and biology 1074: 11–17, 2018.

10. Chao JR, Knight K, Engel AL, Jankowski C, Wang Y, Manson MA, Gu H, Djukovic D, Raftery D, Hurley JB, and Du J. Human retinal pigment epithelial cells prefer proline as a nutrient and transport metabolic intermediates to the retinal side. The Journal of biological chemistry 292: 12895–12905, 2017.

11. Diebold LP, Gil HJ, Gao P, Martinez CA, Weinberg SE, and Chandel NS. Mitochondrial complex III is necessary for endothelial cell proliferation during angiogenesis. Nature metabolism 1: 158–171, 2019.

12. Du J, An J, Linton JD, Wang Y, and Hurley JB. How Excessive cGMP Impacts Metabolic Proteins in Retinas at the Onset of Degeneration. Advances in experimental medicine and biology 1074: 289–295, 2018.

13. Du J, Cleghorn W, Contreras L, Linton JD, Chan GC, Chertov AO, Saheki T, Govindaraju V, Sadilek M, Satrustegui J, and Hurley JB. Cytosolic reducing power preserves glutamate in retina. Proceedings of the National Academy of Sciences of the United States of America 110: 18501–18506, 2013.

14. Du J, Linton JD, and Hurley JB. Probing Metabolism in the Intact Retina Using Stable Isotope Tracers. Methods in enzymology 561: 149–170, 2015.

15. Du J, Rountree A, Cleghorn WM, Contreras L, Lindsay KJ, Sadilek M, Gu H, Djukovic D, Raftery D, Satrustegui J, Kanow M, Chan L, Tsang SH, Sweet IR, and Hurley JB. Phototransduction Influences Metabolic Flux and Nucleotide Metabolism in Mouse Retina. The Journal of biological chemistry 291: 4698–4710, 2016.

16. Du J, Yanagida A, Knight K, Engel AL, Vo AH, Jankowski C, Sadilek M, Tran VT, Manson MA, Ramakrishnan A, Hurley JB, and Chao JR. Reductive carboxylation is a major metabolic pathway in the retinal pigment epithelium. Proceedings of the National Academy of Sciences of the United States of America 113: 14710–14715, 2016.

17. Ehebauer F, Ghavampour S, and Kraus D. Glucose availability regulates nicotinamide N-methyltransferase expression in adipocytes. Life sciences 248: 117474, 2020.

18. Fasullo M, and Endres L. Nucleotide salvage deficiencies, DNA damage and neurodegeneration. International journal of molecular sciences 16: 9431–9449, 2015.

19. Felszeghy S, Viiri J, Paterno JJ, Hyttinen JMT, Koskela A, Chen M, Leinonen H, Tanila H, Kivinen N, Koistinen A, Toropainen E, Amadio M, Smedowski A, Reinisalo M, Winiarczyk M, Mackiewicz J, Mutikainen M, Ruotsalainen AK, Kettunen M, Jokivarsi K, Sinha D, Kinnunen K, Petrovski G, Blasiak J, Bjorkoy G, Koskelainen A, Skottman H, Urtti A, Salminen A, Kannan R, Ferrington DA, Xu H, Levonen AL, Tavi P, Kauppinen A, and Kaarniranta K. Loss of NRF-2 and PGC-1alpha genes leads to retinal pigment epithelium damage resembling dry age-related macular degeneration. Redox biology 20: 1–12, 2019.

20. Ferrington DA, Ebeling MC, Kapphahn RJ, Terluk MR, Fisher CR, Polanco JR, Roehrich H, Leary MM, Geng Z, Dutton JR, and Montezuma SR. Altered bioenergetics and enhanced resistance to oxidative stress in human retinal pigment epithelial cells from donors with age-related macular degeneration. Redox biology 13: 255–265, 2017.

21. Ferrington DA, Fisher CR, and Kowluru RA. Mitochondrial Defects Drive Degenerative Retinal Diseases. Trends in molecular medicine 26: 105–118, 2020.

22. Gibala MJ, Young ME, and Taegtmeyer H. Anaplerosis of the citric acid cycle: role in energy metabolism of heart and skeletal muscle. Acta physiologica Scandinavica 168: 657–665, 2000.

23. Golestaneh N, Chu Y, Cheng SK, Cao H, Poliakov E, and Berinstein DM. Repressed SIRT1/PGC-1alpha pathway and mitochondrial disintegration in iPSC-derived RPE disease model of age-related macular degeneration. Journal of translational medicine 14: 344, 2016.

24. Gong J, Cai H, Noggle S, Paull D, Rizzolo LJ, Del Priore LV, and Fields MA. Stem cell-derived retinal pigment epithelium from patients with age-related macular degeneration exhibit reduced metabolism and matrix interactions. Stem cells translational medicine 9: 364–376, 2020.

25. Grenell A, Wang Y, Yam M, Swarup A, Dilan TL, Hauer A, Linton JD, Philp NJ, Gregor E, Zhu S, Shi Q, Murphy J, Guan T, Lohner D, Kolandaivelu S, Ramamurthy V, Goldberg AFX, Hurley JB, and Du J. Loss of MPC1 reprograms retinal metabolism to impair visual function. Proceedings of the National Academy of Sciences of the United States of America 116: 3530–3535, 2019.

26. Hayaishi O, Nishizuka Y, Tatibana M, Takeshita M, and Kuno S. Enzymatic studies on the metabolism of beta-alanine. The Journal of biological chemistry 236: 781–790, 1961.

27. Hurley JB, Lindsay KJ, and Du J. Glucose, lactate, and shuttling of metabolites in vertebrate retinas. Journal of neuroscience research 93: 1079–1092, 2015.

28. Kanow MA, Giarmarco MM, Jankowski CS, Tsantilas K, Engel AL, D. J, Linton JD, Farnsworth CC, Sloat SR, Rountree A, Sweet IR, Lindsay KJ, Parker ED, Brockerhoff SE, Sadilek M, Chao JR, and Hurley JB. Biochemical adaptations of the retina and retinal pigment epithelium support a metabolic ecosystem in the vertebrate eye. eLife 6: 2017.

29. Kraus D, Yang Q, Kong D, Banks AS, Zhang L, Rodgers JT, Pirinen E, Pulinilkunnil TC, Gong F, Wang YC, Cen Y, Sauve AA, Asara JM, Peroni OD, Monia BP, Bhanot S, Alhonen L, Puigserver P, and Kahn BB. Nicotinamide N-methyltransferase knockdown protects against diet-induced obesity. Nature 508: 258–262, 2014.

30. Kurihara T, Westenskow PD, Gantner ML, Usui Y, Schultz A, Bravo S, Aguilar E, Wittgrove C, Friedlander M, Paris LP, Chew E, Siuzdak G, and Friedlander M. Hypoxia-induced metabolic stress in retinal pigment epithelial cells is sufficient to induce photoreceptor degeneration. eLife 5: 2016.

31. Lai YC, Baker JS, Donti T, Graham BH, Craigen WJ, and Anderson AE. Mitochondrial Dysfunction Mediated by Poly(ADP-Ribose) Polymerase-1 Activation Contributes to Hippocampal Neuronal Damage Following Status Epilepticus. International journal of molecular sciences 18: 2017.

32. Lane AN, and Fan TW. Regulation of mammalian nucleotide metabolism and biosynthesis. Nucleic acids research 43: 2466–2485, 2015.

33. LaVail MM. Rod outer segment disk shedding in rat retina: relationship to cyclic lighting. Science 194: 1071–1074, 1976.

34. Liu L, Su X, Quinn WJ, 3rd, Hui S, Krukenberg K, Frederick DW, Redpath P, Zhan L, Chellappa K, White E, Migaud M, Mitchison TJ, Baur JA, and Rabinowitz JD. Quantitative Analysis of NAD Synthesis-Breakdown Fluxes. Cell metabolism 27: 1067–1080 e1065, 2018.

35. Olsen RKJ, Konarikova E, Giancaspero TA, Mosegaard S, Boczonadi V, Matakovic L, Veauville-Merllie A, Terrile C, Schwarzmayr T, Haack TB, Auranen M, Leone P, Galluccio M, Imbard A, Gutierrez-Rios P, Palmfeldt J, Graf E, Vianey-Saban C, Oppenheim M, Schiff M, Pichard S, Rigal O, Pyle A, Chinnery PF, Konstantopoulou V, Moslinger D, Feichtinger RG, Talim B, Topaloglu H, Coskun T, Gucer S, Botta A, Pegoraro E, Malena A, Vergani L, Mazza D, Zollino M, Ghezzi D, Acquaviva C, Tyni T, Boneh A, Meitinger T, Strom TM, Gregersen N, Mayr JA, Horvath R, Barile M, and Prokisch H. Riboflavin-Responsive and -Non-responsive Mutations in FAD Synthase Cause Multiple Acyl-CoA Dehydrogenase and Combined Respiratory-Chain Deficiency. American journal of human genetics 98: 1130–1145, 2016.

36. Owen OE, Kalhan SC, and Hanson RW. The key role of anaplerosis and cataplerosis for citric acid cycle function. The Journal of biological chemistry 277: 30409–30412, 2002.

37. Phang JM, Liu W, and Zabirnyk O. Proline metabolism and microenvironmental stress. Annual review of nutrition 30: 441–463, 2010.

38. Plana-Bonamaiso A, Lopez-Begines S, Fernandez-Justel D, Junza A, Soler-Tapia A, Andilla J, Loza-Alvarez P, Rosa JL, Miralles E, Casals I, Yanes O, de la Villa P, Buey RM, and Mendez A. Post-translational regulation of retinal IMPDH1 in vivo to adjust GTP synthesis to illumination conditions. eLife 9: 2020.

39. Reyes-Reveles J, Dhingra A, Alexander D, Bragin A, Philp NJ, and Boesze-Battaglia K. Phagocytosis-dependent ketogenesis in retinal pigment epithelium. The Journal of biological chemistry 292: 8038–8047, 2017.

40. Shum M, Houde VP, Bellemare V, Junges Moreira R, Bellmann K, St-Pierre P, Viollet B, Foretz M, and Marette A. Inhibition of mitochondrial complex 1 by the S6K1 inhibitor PF-4708671 partly contributes to its glucose metabolic effects in muscle and liver cells. The Journal of biological chemistry 294: 12250–12260, 2019.

41. Strauss O. The retinal pigment epithelium in visual function. Physiological reviews 85: 845–881, 2005.

42. Sullivan LB, Gui DY, Hosios AM, Bush LN, Freinkman E, and Vander Heiden MG. Supporting Aspartate Biosynthesis Is an Essential Function of Respiration in Proliferating Cells. Cell 162: 552–563, 2015.

43. Tanianskii DA, Jarzebska N, Birkenfeld AL, O’Sullivan JF, and Rodionov RN. Beta-Aminoisobutyric Acid as a Novel Regulator of Carbohydrate and Lipid Metabolism. Nutrients 11: 2019.

44. Titov DV, Cracan V, Goodman RP, Peng J, Grabarek Z, and Mootha VK. Complementation of mitochondrial electron transport chain by manipulation of the NAD+/NADH ratio. Science 352: 231–235, 2016.

45. Wang W, Kini A, Wang Y, Liu T, Chen Y, Vukmanic E, Emery D, Liu Y, Lu X, Jin L, Lee SJ, Scott P, Liu X, Dean K, Lu Q, Fortuny E, James R, Kaplan HJ, D. J, and Dean DC. Metabolic Deregulation of the Blood-Outer Retinal Barrier in Retinitis Pigmentosa. Cell reports 28: 1323–1334 e1324, 2019.

46. Wang Y, Grenell A, Zhong F, Yam M, Hauer A, Gregor E, Zhu S, Lohner D, Zhu J, and Du J. Metabolic signature of the aging eye in mice. Neurobiology of aging 71: 223–233, 2018.

47. Xu R, Ritz BK, Wang Y, Huang J, Zhao C, Gong K, Liu X, and Du J. The retina and retinal pigment epithelium differ in nitrogen metabolism and are metabolically connected. The Journal of biological chemistry 295: 2324–2335, 2020.

48. Yam M, Engel AL, Wang Y, Zhu S, Hauer A, Zhang R, Lohner D, Huang J, Dinterman M, Zhao C, Chao JR, and Du J. Proline mediates metabolic communication between retinal pigment epithelial cells and the retina. The Journal of biological chemistry 294: 10278–10289, 2019.

49. Zabielska MA, Borkowski T, Slominska EM, and Smolenski RT. Inhibition of AMP deaminase as therapeutic target in cardiovascular pathology. Pharmacological reports : PR 67: 682–688, 2015.

50. Zhao C, Yasumura D, Li X, Matthes M, Lloyd M, Nielsen G, Ahern K, Snyder M, Bok D, Dunaief JL, LaVail MM, and Vollrath D. mTOR-mediated dedifferentiation of the retinal pigment epithelium initiates photoreceptor degeneration in mice. The Journal of clinical investigation 121: 369–383, 2011.

