## Supplement Methods for "Inhibition of mitochondrial respiration impairs nutrient consumption and metabolite transport in human retinal pigment epithelium"

Running title: *Metabolic signature of dysfunctional mitochondria*

^1^Department of Ophthalmology, West Virginia University, Morgantown, WV 26506

^2^Department of Biochemistry, West Virginia University, Morgantown, WV 26506

^3^Save Sight Institute, Sydney Medical School, University of Sydney, Sydney, NSW 2000, Australia

^4^Department of Ophthalmology, University of Washington, Seattle, WA 98109
^&^These authors contributed equally to this work.

* Corresponding Authors: Jennifer R. Chao, 750 Republican Street, Box 358058, Seattle WA 98109; Phone: (206) 221-0594;; or Jianhai Du, One Medical Center Dr, PO Box 9193, West Virginia University Eye Institute, Morgantown, WV 26505; Phone: (304)-598-6903; Fax: (304)-598-6928;.

**This PDF file includes:**

Supplemental methods

**Supplemental Methods**

**LDH Activity Assay**

The LDH activity assay was performed using lactate dehydrogenase-SL assay kit (Sekisui Diagnostic, # 327-30 ) according to the manual. Protein, which was extracted from the liver with 50mM K_2_PO_4_ (pH=7.2) served as the positive control. Twenty μl of medium or positive control was mixed with 80μl fresh-made reaction buffer, which contains 260 mM N-methyl-D-glucamine, 40mM lactate, and 2mM NAD. The absorbance at the excitation of 340nm was measured every 30s at 37°C for 20 minutes by the FilterMax F5 Multi-Mode Microplate Reader (Molecular Devices, Sunnyvale, CA, USA). The LDH activity is calculated as delta change, as we previously reported (1, 2) and normalized over the DMSO group.

Reference

1.	**Du J, Yanagida A, Knight K, Engel AL, Vo AH, Jankowski C, Sadilek M, Tran VT, Manson MA, Ramakrishnan A, Hurley JB, and Chao JR**. Reductive carboxylation is a major metabolic pathway in the retinal pigment epithelium. *Proceedings of the National Academy of Sciences of the United States of America* 113: 14710-14715, 2016.

2.	**Yam M, Engel AL, Wang Y, Zhu S, Hauer A, Zhang R, Lohner D, Huang J, Dinterman M, Zhao C, Chao JR, and Du J**. Proline mediates metabolic communication between retinal pigment epithelial cells and the retina. *The Journal of biological chemistry* 294: 10278-10289, 2019.
