## Supplement Tables for "Inhibition of mitochondrial respiration impairs nutrient consumption and metabolite transport in human retinal pigment epithelium"

Running title: *Metabolic signature of dysfunctional mitochondria*

^1^Department of Ophthalmology, West Virginia University, Morgantown, WV 26506

^2^Department of Biochemistry, West Virginia University, Morgantown, WV 26506

^3^Save Sight Institute, Sydney Medical School, University of Sydney, Sydney, NSW 2000, Australia

^4^Department of Ophthalmology, University of Washington, Seattle, WA 98109
^&^These authors contributed equally to this work.

* Corresponding Authors: Jennifer R. Chao, 750 Republican Street, Box 358058, Seattle WA 98109; Phone: (206) 221-0594;; or Jianhai Du, One Medical Center Dr, PO Box 9193, West Virginia University Eye Institute, Morgantown, WV 26505; Phone: (304)-598-6903; Fax: (304)-598-6928;.

**This PDF file includes:**

Table S1-S6

**Table S1**. Significantly changed metabolites in 1h media after inhibition of mitochondrial respiration.

| **Metabolite** | **f value** | **p value** | **FDR** |
| --- | --- | --- | --- |
| Lactate | 84.056 | 2.17E-06 | 0.000187 |
| Cis-aconitic acid | 18.863 | 0.00055 | 0.018053 |
| Glycerol | 15.177 | 0.001152 | 0.018053 |
| oxalic acid | 14.905 | 0.001223 | 0.018053 |
| 3-Aminoisobutanoic acid | 14.088 | 0.001475 | 0.018053 |
| N1-Methylnicotinamide | 13.916 | 0.001536 | 0.018053 |
| Glucose | 13.744 | 0.0016 | 0.018053 |
| Uracil | 13.543 | 0.001679 | 0.018053 |
| Succinate | 10.919 | 0.003354 | 0.032047 |
| 4-hydroxyproline | 10.388 | 0.003918 | 0.033697 |
| Proline | 9.3191 | 0.005466 | 0.042732 |

Data were analyzed with One Way-Anova using Fisher’s FSD by MetaboAnalyst 4.0. False discovery rate (FDR).

**Table S2**. Significantly changed metabolites in 6h media after inhibition of mitochondrial respiration.

| **Metabolite** | **f value** | **p value** | **FDR** |
| --- | --- | --- | --- |
| Lactate | 1183.4 | 6.28E-11 | 5.53E-09 |
| Glucose | 957.35 | 1.46E-10 | 6.44E-09 |
| N-Aetyl-Glycine | 658.48 | 6.50E-10 | 1.91E-08 |
| Proline | 143.75 | 2.69E-07 | 5.91E-06 |
| Succinate | 118.25 | 5.77E-07 | 1.02E-05 |
| Choline | 76.951 | 3.06E-06 | 4.48E-05 |
| 3-Aminoisobutanoic acid | 56.382 | 1.00E-05 | 0.000126 |
| 3-Hydroxybutyric acid | 53.695 | 1.21E-05 | 0.000127 |
| Isoleucine | 52.669 | 1.30E-05 | 0.000127 |
| Leucine | 44.399 | 2.48E-05 | 0.000201 |
| N-Asp | 44.257 | 2.51E-05 | 0.000201 |
| N1-Methylnicotinamide | 40.625 | 3.45E-05 | 0.000253 |
| a-Ketoglutarate | 28.659 | 0.000125 | 0.000806 |
| Hypoxanthine | 28.441 | 0.000128 | 0.000806 |
| Pyruvate | 27.466 | 0.000146 | 0.000854 |
| Cis-aconitic acid | 25.106 | 0.000201 | 0.001105 |
| Uracil | 21.365 | 0.000356 | 0.001844 |
| biotin | 20.956 | 0.000381 | 0.001863 |
| Cytidine | 15.79 | 0.001008 | 0.00467 |
| Valine | 13.315 | 0.001776 | 0.007813 |
| Citrate | 12.257 | 0.002322 | 0.009563 |
| Isocitrate | 12.147 | 0.002391 | 0.009563 |
| Erythritol | 11.679 | 0.00271 | 0.01037 |
| Glycine | 11.105 | 0.00318 | 0.011659 |
| Betaine | 10.725 | 0.003547 | 0.012486 |
| Glutamine | 10.39 | 0.003916 | 0.013253 |
| GAP | 9.5493 | 0.005076 | 0.016543 |
| hypotaurine | 9.3035 | 0.005494 | 0.017265 |
| Inosine | 8.7804 | 0.006536 | 0.019833 |
| Methionine | 8.6768 | 0.006771 | 0.019834 |
| Phenylpyruvate | 8.5853 | 0.006987 | 0.019834 |
| Uridine | 7.534 | 0.010218 | 0.028099 |

**Table S3**. Significantly changed metabolites in 24h media after inhibition of mitochondrial respiration.

| **Metabolite** | **f value** | **p value** | **FDR** |
| --- | --- | --- | --- |
| N-Aetyl-Glycine | 396.83 | 4.87E-09 | 4.34E-07 |
| Citrate | 194.91 | 8.12E-08 | 2.72E-06 |
| Isocitrate | 189.02 | 9.17E-08 | 2.72E-06 |
| Guanine | 81.15 | 2.49E-06 | 3.75E-05 |
| Glucose | 80.943 | 2.51E-06 | 3.75E-05 |
| Proline | 80.841 | 2.53E-06 | 3.75E-05 |
| Butyrylcarnitine | 51.732 | 1.39E-05 | 0.000177 |
| 3-Aminoisobutanoic acid | 47.674 | 1.90E-05 | 0.000206 |
| Phenylpyruvate | 45.577 | 2.25E-05 | 0.000206 |
| Propionylcarnitine | 44.972 | 2.36E-05 | 0.000206 |
| Choline | 44.091 | 2.54E-05 | 0.000206 |
| Hypoxanthine | 42.447 | 2.93E-05 | 0.000217 |
| Erythritol | 38.493 | 4.22E-05 | 0.000279 |
| 3-Hydroxybutyric acid | 38.096 | 4.39E-05 | 0.000279 |
| hypotaurine | 35.926 | 5.45E-05 | 0.000324 |
| Guanosine | 35.298 | 5.82E-05 | 0.000324 |
| Nicotinamide | 32.8 | 7.62E-05 | 0.000399 |
| Succinate | 32.115 | 8.24E-05 | 0.000407 |
| Uracil | 30.207 | 0.000103 | 0.000483 |
| N1-Methylnicotinamide | 26.107 | 0.000175 | 0.000777 |
| Citrulline | 24.348 | 0.000224 | 0.00095 |
| Cytidine | 22.645 | 0.00029 | 0.001126 |
| Isobutyrylcarnitine | 22.627 | 0.000291 | 0.001126 |
| Betaine | 20.094 | 0.000441 | 0.001637 |
| Valine | 15.654 | 0.001038 | 0.003696 |
| Creatine | 14.603 | 0.00131 | 0.004483 |
| Alanine | 13.944 | 0.001526 | 0.00503 |
| Cis-aconitic acid | 13.213 | 0.001821 | 0.005788 |
| N-Asp | 12.666 | 0.002089 | 0.006411 |
| myo inositol | 12.411 | 0.002231 | 0.006618 |
| Pyruvate | 12.14 | 0.002395 | 0.006877 |
| Ethanolamine | 10.564 | 0.003718 | 0.010341 |
| Leucine | 9.4832 | 0.005184 | 0.013981 |
| PEP | 9.2877 | 0.005522 | 0.014313 |
| Isoleucine | 9.229 | 0.005629 | 0.014313 |
| Glutamine | 8.9763 | 0.006119 | 0.014874 |
| IPP | 8.9447 | 0.006184 | 0.014874 |
| Xanthine | 7.5011 | 0.010346 | 0.024232 |
| Acetyl L Carnitine | 7.3605 | 0.010918 | 0.024916 |
| Urea | 7.2707 | 0.011304 | 0.025152 |
| Lactate | 6.8862 | 0.013164 | 0.028575 |
| a-Ketoglutarate | 6.2399 | 0.017237 | 0.036525 |
| Inosine | 5.6898 | 0.02201 | 0.045556 |

**Table S4**. Significantly changed metabolites in cells at 24h after inhibition of mitochondrial respiration.

| **Metabolite** | **f value** | **p value** | **FDR** |
| --- | --- | --- | --- |
| Phosphocreatine | 7112.1 | 4.86E-14 | 6.12E-12 |
| 2-Methylbutyroylcarnitine | 2075.3 | 6.67E-12 | 3.11E-10 |
| AMP | 2021.9 | 7.40E-12 | 3.11E-10 |
| Isobutyrylcarnitine | 720.15 | 4.55E-10 | 1.34E-08 |
| Oxidized glutathione | 671.69 | 6.01E-10 | 1.34E-08 |
| Butyrylcarnitine | 661.07 | 6.40E-10 | 1.34E-08 |
| Myristoylcarnitine | 567.3 | 1.18E-09 | 2.12E-08 |
| Glucose | 543.96 | 1.39E-09 | 2.19E-08 |
| IMP | 501.9 | 1.92E-09 | 2.55E-08 |
| Aconitate | 495.14 | 2.02E-09 | 2.55E-08 |
| NADH | 446.09 | 3.06E-09 | 3.51E-08 |
| Citraconic Acid | 436.14 | 3.35E-09 | 3.52E-08 |
| Succinate | 282.87 | 1.87E-08 | 1.81E-07 |
| 3-HB | 267.34 | 2.33E-08 | 2.10E-07 |
| IPP | 258.01 | 2.68E-08 | 2.25E-07 |
| 2-hydroxyglutarate | 233.05 | 4.01E-08 | 3.16E-07 |
| Myo inositol | 201.47 | 7.13E-08 | 5.28E-07 |
| 4-hydroxyproline | 184.59 | 1.01E-07 | 7.04E-07 |
| a-Ketoglutarate | 161.55 | 1.70E-07 | 1.13E-06 |
| Propionylcarnitine | 153.46 | 2.08E-07 | 1.25E-06 |
| CAMP | 153.31 | 2.09E-07 | 1.25E-06 |
| Hypotaurine | 151.49 | 2.19E-07 | 1.25E-06 |
| Citrulline | 144.62 | 2.62E-07 | 1.38E-06 |
| N-Aetyl-Glycine | 144.57 | 2.63E-07 | 1.38E-06 |
| NADPH | 115.9 | 6.24E-07 | 3.01E-06 |
| Isocitrate | 115.06 | 6.42E-07 | 3.01E-06 |
| NADP | 114.97 | 6.44E-07 | 3.01E-06 |
| Betaine | 110.98 | 7.39E-07 | 3.22E-06 |
| Citrate | 110.85 | 7.42E-07 | 3.22E-06 |
| NAD | 109.62 | 7.75E-07 | 3.26E-06 |
| Cytosine | 92.405 | 1.51E-06 | 6.12E-06 |
| 1-Methyladenosine | 89.12 | 1.73E-06 | 6.82E-06 |
| Adenine | 86.282 | 1.96E-06 | 7.50E-06 |
| Uracil | 81.54 | 2.44E-06 | 9.06E-06 |
| Argininosuccinate | 74.84 | 3.40E-06 | 1.19E-05 |
| Hypoxanthine | 74.826 | 3.40E-06 | 1.19E-05 |
| Carnosine | 73.718 | 3.60E-06 | 1.21E-05 |
| Lysine | 73.558 | 3.63E-06 | 1.21E-05 |
| PEP | 68.663 | 4.73E-06 | 1.53E-05 |
| Choline | 60.793 | 7.54E-06 | 2.38E-05 |
| UDP | 55.99 | 1.03E-05 | 3.17E-05 |
| Glucose 1-phosphate | 54.489 | 1.14E-05 | 3.43E-05 |
| Aminoadipic acid | 53.834 | 1.20E-05 | 3.46E-05 |
| Cytidine | 53.728 | 1.21E-05 | 3.46E-05 |
| 3PG | 53.143 | 1.26E-05 | 3.52E-05 |
| Taurine | 51.152 | 1.45E-05 | 3.98E-05 |
| 3-Aminoisobutanoic acid | 50.463 | 1.53E-05 | 4.10E-05 |
| Proline | 50.08 | 1.57E-05 | 4.11E-05 |
| Ornithine | 49.898 | 1.60E-05 | 4.11E-05 |
| N-Asp | 48.631 | 1.76E-05 | 4.35E-05 |
| Gamma-Aminobutyric acid | 48.615 | 1.76E-05 | 4.35E-05 |
| Guanine | 47.679 | 1.90E-05 | 4.51E-05 |
| Hexanoylcarnitine | 47.666 | 1.90E-05 | 4.51E-05 |
| Leucine | 47.091 | 1.99E-05 | 4.57E-05 |
| ADP | 47.05 | 1.99E-05 | 4.57E-05 |
| Isoleucine | 44.614 | 2.43E-05 | 5.48E-05 |
| Guanosine | 43.517 | 2.67E-05 | 5.91E-05 |
| Histamine | 42.402 | 2.94E-05 | 6.40E-05 |
| Malate | 41.407 | 3.22E-05 | 6.87E-05 |
| Pyroglutamic acid | 39.511 | 3.83E-05 | 8.05E-05 |
| Glycerol | 39.216 | 3.94E-05 | 8.14E-05 |
| Thiamine | 38.175 | 4.35E-05 | 8.85E-05 |
| Pantothenic acid | 37.817 | 4.51E-05 | 8.94E-05 |
| Glucose 6-phosphate | 37.75 | 4.54E-05 | 8.94E-05 |
| Lactate | 36.004 | 5.41E-05 | 0.000105 |
| Valine | 33.06 | 7.41E-05 | 0.000141 |
| ATP | 30.693 | 9.72E-05 | 0.000183 |
| Glycerate | 29.992 | 0.000106 | 0.000196 |
| Aspartate | 28.177 | 0.000133 | 0.000242 |
| GAP | 27.301 | 0.000149 | 0.000268 |
| Acetylglycine | 25.91 | 0.00018 | 0.000319 |
| Xanthine | 24.614 | 0.000216 | 0.000377 |
| Glutamine | 23.262 | 0.000264 | 0.000452 |
| Acetyl L Carnitine | 23.216 | 0.000266 | 0.000452 |
| biotin | 22.588 | 0.000293 | 0.000492 |
| Glutaric Acid | 21.46 | 0.000351 | 0.000581 |
| Erythritol | 20.339 | 0.000423 | 0.000692 |
| Glycine | 20.238 | 0.000431 | 0.000695 |
| Methionine | 19.516 | 0.000489 | 0.000779 |
| L-Arginine | 18.662 | 0.00057 | 0.000899 |
| Trigonelline | 17.496 | 0.000712 | 0.001108 |
| Tryptophan | 16.487 | 0.000872 | 0.001339 |
| Nicotinamide | 16.214 | 0.000922 | 0.001386 |
| Urea | 16.205 | 0.000924 | 0.001386 |
| N1-Methylnicotinamide | 14.959 | 0.001209 | 0.001792 |
| Palmitate | 14.851 | 0.001238 | 0.001815 |
| Acetoacetate | 14.43 | 0.001363 | 0.001973 |
| Phenyalanine | 14.363 | 0.001384 | 0.001981 |
| Stearic acid | 13.634 | 0.001643 | 0.002326 |
| Creatine | 13.554 | 0.001675 | 0.002345 |
| Xanthurenic acid | 13.446 | 0.00172 | 0.002381 |
| Histidine | 13.343 | 0.001764 | 0.002415 |
| Cystathionine | 13.198 | 0.001827 | 0.002461 |
| Erythrono 1,4 Lactone | 13.179 | 0.001836 | 0.002461 |
| Maleic Acid | 12.136 | 0.002398 | 0.00318 |
| Asparagene | 11.676 | 0.002713 | 0.00356 |
| Serine | 11.552 | 0.002806 | 0.003645 |
| Hippurate | 11.456 | 0.002882 | 0.003706 |
| Adipic Acid | 11.38 | 0.002943 | 0.003746 |
| Fumarate | 11.056 | 0.003225 | 0.004063 |
| Cis-aconitic acid | 10.878 | 0.003394 | 0.004234 |
| Creatinine | 10.382 | 0.003925 | 0.004849 |
| Oxalic acid | 10.345 | 0.003969 | 0.004856 |
| L-Asparagine | 10.013 | 0.00439 | 0.005319 |
| D-Ribulose 5-phosphate | 9.9053 | 0.004538 | 0.005446 |
| N-alpha-Acetyl-L-lysine | 9.628 | 0.00495 | 0.005884 |
| Threonine | 9.2331 | 0.005621 | 0.006619 |
| Succinic semiadehyde | 8.9561 | 0.00616 | 0.007187 |
| L-Homoserine | 8.8324 | 0.006422 | 0.007423 |
| Riboflavin | 8.3701 | 0.007531 | 0.008626 |
| Glutathione | 7.8015 | 0.009244 | 0.010493 |
| Tyrosine | 7.2492 | 0.0114 | 0.012824 |
| Urate | 6.9685 | 0.012735 | 0.014201 |
| Alanine | 6.8475 | 0.013371 | 0.014779 |
| Beta-Alanine | 6.5583 | 0.015059 | 0.0165 |
| Glutamate | 6.5207 | 0.015298 | 0.016617 |
| Cystine | 4.4229 | 0.04115 | 0.044315 |
| carbonate | 4.3858 | 0.041974 | 0.044819 |

**Table S5**. **The reagents and key resources.**

| **Reagent or Media components** | **SOURCE** | **IDENTIFIER** |
| --- | --- | --- |
| Piericidin | Cayman Chemical | 15379 |
| Antimycin | Sigma-Aldrich | A8674 |
| Oligomycin | Sigma-Aldrich | 75351 |
| DMSO | Sigma-Aldrich | D2650 |
| [^13^C_6_]-Glucose | Cambridge-Isotope Laboratories | CLM-1396 |
| MEM a | Fisher | 12561056 |
| Non-essential amino acids | Life Technologies (Invitrogen) | 1140-050 |
| N1 medium supplement | Sigma-Aldrich | N6530-5mL |
| FBS | Atlanta Biologicals | S11550 |
| Taurine | Sigma-Aldrich | T0625-10G |
| Hydrocortisone | Sigma-Aldrich | H0396-100MG |
| 3, 3', 5-Triiodo-L-Thyronine | Sigma-Aldrich | T6397-100MG |
| Penicillin-Streptomycin (5,000 U/mL) | Life Technologies (Invitrogen) | 15070-063 |
| DMEM, no glucose, no glutamine, no phenol red | ThermoFisher | A1443001 |
| Y-27632 dihydrochloride | Tocris/R&D Systems | 1254 |
| **Cytotoxicity experiments** |  |  |
| Lactate Dehydrogenase-SL (kit) | Sekisui Diagnostics | 327-30 |
| **Mass Spectrometry (LC MS and GC MS)** |  |  |
| Methanol, Optima™ LC/MS Grade | Fisher Chemical | A456-500 |
| Acquity UPLC BEH Amide 1.7 µm Vanguard pre-column 2.1 x 5mm column | Waters | 186004799 |
| Acquity UPLC BEH Amide 1.7 µm 2.1 x 50mm column | Waters | 186004800 |
| Water, Optima™ LC/MS Grade | Fisher Chemical | 7732-18-5 |
| Acetonitrile, Optima™ LC/MS Grade | Fisher Chemical | 75-05-8 |
| Ammonium Acetate | Sigma-Aldrich | 431311 |
| Ammonium hydroxide 28 - 30% in water ACS | VWR | AC42330-5000 |
| Methoxyamine hydrochloride | Sigma-Aldrich | 226904 |
| Pyridine | Sigma-Aldrich | 270970 |
| N-tert-Butyldimethylsilyl-N-methyltrifluoroacetamide | Sigma-Aldrich | 394882 |
| DB-5ms GC Column, 30 m, 0.25 mm, 0.25 µm, | Agilent Technologies | 122-5532 |

**Table S6.** The list of metabolites and parameters for LC MS and GC MS

| **Names** | **Nutrients** | **Precursor (Da)** | **Product (Da)** | **Declustering potential** | **Collision Energy** | **HMDB** | **Platform** | **Polarity** |
| --- | --- | --- | --- | --- | --- | --- | --- | --- |
| Acetylglycine | Amino acid | 116.0 | 74.0 | -37.0 | -14.0 | HMDB00532 | LCMS | - |
| Aconitate | Glucose | 173.0 | 85.0 | -37.0 | -18.0 | HMDB00072 | LCMS | - |
| Adenine | Nucleotide | 134.0 | 107.1 | -92.0 | -25.0 | HMDB00034 | LCMS | - |
| Adenosine 5'-diphosphate | Energy | 426.0 | 79.1 | -65.0 | -84.0 | HMDB01341 | LCMS | - |
| Aminoadipic acid | Amino acid | 160.1 | 116.1 | -47.0 | -20.0 | HMDB00510 | LCMS | - |
| Adenosine monophosphate | Energy | 346.0 | 134.0 | -82.0 | -40.0 | HMDB00045 | LCMS | - |
| Cyclic AMP | Nucleotide | 328.0 | 134.0 | -87.0 | -32.0 | HMDB00058 | LCMS | - |
| Citraconic Acid | Glucose | 129.0 | 85.0 | -21.0 | -12.0 | HMDB00634 | LCMS | - |
| Citrulline | Amino acid | 174.1 | 131.1 | -32.0 | -20.0 | HMDB00904 | LCMS | - |
| Creatinine | Amino acid | 112.0 | 41.0 | -67.0 | -36.0 | HMDB00562 | LCMS | - |
| D-Ribulose 5-phosphate | Nucleotide | 229.0 | 79.0 | -70.0 | -45.0 | HMDB0000618 | LCMS | - |
| Glucose | Glucose | 179.0 | 89.0 | -50.0 | -15.0 | HMDB00122 | LCMS | - |
| Glucose 1-phosphate | Glucose | 259.0 | 79.0 | -85.0 | -60.0 | HMDB0001586 | LCMS | - |
| Glucose 6-phosphate | Glucose | 259.0 | 79.0 | -46.0 | -62.0 | HMDB01401 | LCMS | - |
| Glutaric Acid | Amino acid | 131.0 | 87.0 | -45.0 | -16.0 | HMDB00661 | LCMS | - |
| Glutathione | Energy | 306.0 | 143.1 | -61.0 | -26.0 | HMDB00125 | LCMS | - |
| Guanosine | Nucleotide | 282.1 | 150.0 | -67.0 | -33.0 | HMDB00133 | LCMS | - |
| Hippurate | Amino acid | 178.0 | 77.1 | -61.0 | -23.0 | HMDB00714 | LCMS | - |
| Hypotaurine | Amino acid | 108.1 | 64.0 | -40.0 | -17.0 | HMDB0000965 | LCMS | - |
| Hypoxanthine | Nucleotide | 135.0 | 65.0 | -108.0 | -37.0 | HMDB00157 | LCMS | - |
| Inosinic acid | Nucleotide | 347.0 | 79.0 | -118.0 | -86.0 | HMDB00175 | LCMS | - |
| Inosine | Nucleotide | 267.0 | 135.0 | -123.0 | -30.0 | HMDB0000195 | LCMS | - |
| Myo inositol | Amino acid | 179.0 | 87.0 | -105.0 | -24.0 | HMDB0000211 | LCMS | - |
| NADH | NAD metabolism | 664.0 | 397.0 | -7.0 | -46.0 | HMDB01487 | LCMS | - |
| Oxalic acid | Nucleotide | 89.0 | 61.0 | -45.0 | -10.0 | HMDB0002329 | LCMS | - |
| Pantothenic acid | Vitamin | 218.1 | 71.0 | -70.0 | -41.0 | HMDB0000210 | LCMS | - |
| Phosphocreatine | Amino acid | 210.0 | 79.0 | -42.0 | -49.0 | HMDB0041624 | LCMS | - |
| Uridine 5'-diphosphate | Nucleotide | 403.0 | 79.0 | -30.0 | -73.0 | HMDB00295 | LCMS | - |
| Urate | Nucleotide | 167.0 | 124.0 | -86.0 | -19.0 | HMDB0000289 | LCMS | - |
| Xanthine | Nucleotide | 151.0 | 108.0 | -70.0 | -23.0 | HMDB00292 | LCMS | - |
| Xanthurenic acid | Nucleotide | 204.0 | 160.0 | -67.0 | -19.0 | HMDB00881 | LCMS | - |
| Adipic Acid | Dicarboxylic acids | 145.0 | 83.0 | -50.0 | -17.0 | HMDB0000448 | LCMS | - |
| Isopentenyl pyrophosphate | Lipid | 245.1 | 79.0 | -60.0 | -50.0 | HMDB0001347 | LCMS | - |
| 1-Methyladenosine | Nucleotide | 282.1 | 150.1 | 48.0 | 35.0 | HMDB03331 | LCMS | + |
| 3-Aminoisobutanoic acid | Nucleotide | 104.1 | 58.0 | 55.0 | 38.0 | HMDB03911 | LCMS | + |
| Acetyl L Carnitine | Carnitine | 204.1 | 85.0 | 71.0 | 30.0 | HMDB00201 | LCMS | + |
| Adenosine | Nucleotide | 268.1 | 136.0 | 95.0 | 38.0 | HMDB00050 | LCMS | + |
| Alanine | Amino acid | 90.0 | 44.0 | 50.0 | 20.0 | HMDB0000161 | LCMS | + |
| ATP | Energy | 507.9 | 136.1 | 14.0 | 60.0 | HMDB00538 | LCMS | + |
| Betaine | Amino acid | 118.1 | 58.0 | 166.0 | 56.0 | HMDB00043 | LCMS | + |
| Carnosine | Amino acid | 227.1 | 110.1 | 157.0 | 32.0 | HMDB00033 | LCMS | + |
| Choline | Amino acid | 104.1 | 60.1 | 95.0 | 37.0 | HMDB00097 | LCMS | + |
| Creatine | Amino acid | 132.1 | 90.0 | 170.0 | 17.0 | HMDB00064 | LCMS | + |
| Cytidine | Nucleotide | 244.0 | 112.1 | 50.0 | 30.0 | HMDB00089 | LCMS | + |
| Cytosine | Nucleotide | 112.0 | 95.0 | 97.0 | 23.0 | HMDB0000630 | LCMS | + |
| Flavin adenine diNucleotide | Energy | 786.1 | 136.1 | 52.0 | 43.0 | HMDB01248 | LCMS | + |
| Glutamine | Amino acid | 147.1 | 84.1 | 45.0 | 23.0 | HMDB00641 | LCMS | + |
| L-Arginine | Amino acid | 175.1 | 70.1 | 45.0 | 33.0 | HMDB00517 | LCMS | + |
| L-Asparagine | Amino acid | 133.1 | 70.1 | 55.0 | 24.0 | HMDB0033780 | LCMS | + |
| L-Homoserine | Amino acid | 120.1 | 74.0 | 37.0 | 16.0 | HMDB00719 | LCMS | + |
| Lysine | Amino acid | 147.1 | 84.0 | 38.0 | 22.0 | HMDB0000182 | LCMS | + |
| 1-Methylnicotinamide | Nucleotide | 137.0 | 78.0 | 110.0 | 37.0 | HMDB00699 | LCMS | + |
| NAD | NAD metabolism | 664.0 | 136.0 | 27.0 | 48.0 | HMDB00902 | LCMS | + |
| NADP | NAD metabolism | 744.0 | 136.0 | 9.0 | 78.0 | HMDB00217 | LCMS | + |
| NADPH | NAD metabolism | 746.0 | 302.0 | 20.0 | 44.0 | HMDB00221 | LCMS | + |
| Nicotinamide | NAD metabolism | 123.0 | 80.0 | 111.0 | 20.0 | HMDB01406 | LCMS | + |
| Ornithine | Amino acid | 133.0 | 70.0 | 47.0 | 23.0 | HMDB0000214 | LCMS | + |
| Oxidized glutathione | Energy | 613.0 | 231.0 | 16.0 | 42.0 | HMDB03337 | LCMS | + |
| Proline | Amino acid | 116.1 | 70.1 | 70.0 | 44.0 | HMDB00162 | LCMS | + |
| Pyroglutamic acid | Amino acid | 130.1 | 84.0 | 100.0 | 20.0 | HMDB0000267 | LCMS | + |
| Riboflavin | Nucleotide | 377.1 | 243.1 | 14.0 | 29.0 | HMDB00244 | LCMS | + |
| Serine | Amino acid | 106.0 | 60.0 | 40.0 | 22.0 | HMDB0000187 | LCMS | + |
| Thiamine | Vitamin | 265.0 | 122.1 | 67.0 | 40.0 | HMDB00235 | LCMS | + |
| Threonine | Amino acid | 120.1 | 102.0 | 50.0 | 10.0 | HMDB0000167 | LCMS | + |
| Trigonelline | NAD metabolism | 138.0 | 92.0 | 60.0 | 27.0 | HMDB00875 | LCMS | + |
| Tryptophan | Amino acid | 205.0 | 146.0 | 75.0 | 35.0 | HMDB00929 | LCMS | + |
| Tyrosine | Amino acid | 182.1 | 136.0 | 40.0 | 17.0 | HMDB0000158 | LCMS | + |
| Valine | Amino acid | 118.1 | 72.0 | 60.0 | 14.0 | HMDB0000883 | LCMS | + |
| 2-MethylbutyroylCarnitine | Carnitine | 246.2 | 85.1 | 100.0 | 43.0 | HMDB0000378 | LCMS | + |
| ButyrylCarnitine | Carnitine | 232.2 | 85.1 | 72.0 | 30.0 | HMDB0002013 |  | + |
| Gamma-Aminobutyric acid | Lipid | 104.1 | 87.0 | 57.0 | 13.0 | HMDB0000112 | LCMS | + |
| Guanine | Nucleotide | 152.0 | 110.0 | 20.0 | 28.0 | HMDB0000132 | LCMS | + |
| HexanoylCarnitine | Carnitine | 260.2 | 85.1 | 80.0 | 25.0 | HMDB0000705 | LCMS | + |
| Histamine | Amino acid | 112.0 | 95.0 | 65.0 | 18.0 | HMDB0000870 | LCMS | + |
| IsobutyrylCarnitine | Carnitine | 232.1 | 85.1 | 70.0 | 30.0 | HMDB0000736 | LCMS | + |
| L-PalmitoylCarnitine | Carnitine | 400.4 | 85.1 | 30.0 | 40.0 | HMDB0000222 | LCMS | + |
| PropionylCarnitine | Carnitine | 218.1 | 85.1 | 60.0 | 24.0 | HMDB0000824 | LCMS | + |
| Argininosuccinate | Amino acid | 291.1 | 69.9 | 40.0 | 32.0 | HMDB0000052 | LCMS | + |
| Cystathionine | Amino acid | 221.1 | 79.1 | -87.0 | -30.0 | HMDB0000099 | LCMS | - |
| Erythritol | Sugar | 123.0 | 80.0 | 40.0 | 33.0 | HMDB0002994 | LCMS | + |
| MyristoylCarnitine | Carnitine | 372.4 | 85.1 | 70.0 | 25.0 | HMDB0005066 | LCMS | + |
| N-alpha-Acetyl-L-lysine | Amino acid | 189.1 | 84.1 | 50.0 | 32.0 | HMDB0000446 | LCMS | + |
| 2-hydroxyglutarate | Glucose | 433.2 | NA | NA | NA | HMDB0059655 | GCMS | + |
| 3-Hydroxybutyric acid | Ketone body | 275.1 | NA | NA | NA | HMDB0000357 | GCMS | + |
| 3-Phosphoglyceric acid | Glucose | 585.3 | NA | NA | NA | HMDB0000807 | GCMS | + |
| 4-hydroxyproline | Amino acid | 314.2 | NA | NA | NA | HMDB00725 | GCMS | + |
| Acetoacetate | Ketone body | 188.1 | NA | NA | NA | HMDB0000060 | GCMS | + |
| a-Ketoglutarate | Glucose | 346.0 | NA | NA | NA | HMDB0000208 | GCMS | + |
| Asparagene | Amino acid | 417.0 | NA | NA | NA | HMDB0000168 | GCMS | + |
| Aspartate | Amino acid | 418.0 | NA | NA | NA | HMDB0000191 | GCMS | + |
| Cholesterol | Lipid | 443.4 | NA | NA | NA | HMDB0000067 | GCMS | + |
| Citrate | Glucose | 459.0 | NA | NA | NA | HMDB0000094 | GCMS | + |
| Cysteine | Amino acid | 304.2 | NA | NA | NA | HMDB0000574 | GCMS | + |
| Cystine | Amino acid | 302.2 | NA | NA | NA | HMDB0000192 | GCMS | + |
| Fumarate | Glucose | 287.0 | NA | NA | NA | HMDB0000134 | GCMS | + |
| Glutamate | Amino acid | 432.0 | NA | NA | NA | HMDB0003339 | GCMS | + |
| Glycerate | Lipid | 391.2 | NA | NA | NA | HMDB0000139 | GCMS | + |
| Glycine | Amino acid | 218.0 | NA | NA | NA | HMDB0000123 | GCMS | + |
| Histidine | Amino acid | 196.1 | NA | NA | NA | HMDB0000177 | GCMS | + |
| Isocitrate | Glucose | 591.3 | NA | NA | NA | HMDB0000193 | GCMS | + |
| Isoleucine | Amino acid | 200.0 | NA | NA | NA | HMDB0000172 | GCMS | + |
| Lactate | Glucose | 261.0 | NA | NA | NA | HMDB0000190 | GCMS | + |
| Leucine | Amino acid | 200.0 | NA | NA | NA | HMDB0000687 | GCMS | + |
| Malate | Glucose | 419.2 | NA | NA | NA | HMDB0031518 | GCMS | + |
| Methionine | Amino acid | 218.0 | NA | NA | NA | HMDB0000696 | GCMS | + |
| Palmitate | Lipid | 313.2 | NA | NA | NA | HMDB0000220 | GCMS | + |
| Phosphoenolpyruvic acid | Glucose | 453.1 | NA | NA | NA | HMDB0000263 | GCMS | + |
| Phenylalanine | Amino acid | 302.0 | NA | NA | NA | HMDB0000159 | GCMS | + |
| Pyruvate | Glucose | 174.0 | NA | NA | NA | HMDB0000243 | GCMS | + |
| Stearic acid | Lipid | 341.3 | NA | NA | NA | HMDB0000827 | GCMS | + |
| Succinate | Glucose | 289.0 | NA | NA | NA | HMDB0000254 | GCMS | + |
| Succinic semiadehyde | Amino acid | 159.1 | NA | NA | NA | HMDB0001259 | GCMS | + |
| Taurine | Amino acid | 296.0 | NA | NA | NA | HMDB0000251 | GCMS | + |
| Uracil | Nucleotide | 283.3 | NA | NA | NA | HMDB00300 | GCMS | + |
| Urea | Nucleotide | 231.1 | NA | NA | NA | HMDB0000294 | GCMS | + |
| Beta-Alanine | Amino acid | 260.2 | NA | NA | NA | HMDB0000056 | GCMS | + |
| Biotin | Vitamin | 529.4 | NA | NA | NA | HMDB0000030 | GCMS | + |
| Carbonate | Glucose | 233.1 | NA | NA | NA | HMDB0031453 | GCMS | + |
| Cis-aconitic acid | Glucose | 459.2 | NA | NA | NA | HMDB0000072 | GCMS | + |
| Glyceraldehyde 3-phosphate | Glucose | 484.2 | NA | NA | NA | HMDB0001112 | GCMS | + |
| Glycerol | Lipid | 377.2 | NA | NA | NA | HMDB0000131 | GCMS | + |
| Maleic Acid | Glucose | 287.1 | NA | NA | NA | HMDB0000176 | GCMS | + |
| N-Acetylglycine | Amino acid | 288.1 | NA | NA | NA | HMDB0000532 | GCMS | + |
| N-Acetyl-L-aspartic acid | Amino acid | 460.2 | NA | NA | NA | HMDB0000812 | GCMS | + |
