## Supplement Figures for "Inhibition of mitochondrial respiration impairs nutrient consumption and metabolite transport in human retinal pigment epithelium"

Running title: *Metabolic signature of dysfunctional mitochondria*

^1^Department of Ophthalmology, West Virginia University, Morgantown, WV 26506

^2^Department of Biochemistry, West Virginia University, Morgantown, WV 26506

^3^Save Sight Institute, Sydney Medical School, University of Sydney, Sydney, NSW 2000, Australia

^4^Department of Ophthalmology, University of Washington, Seattle, WA 98109
^&^These authors contributed equally to this work.

* Corresponding Authors: Jennifer R. Chao, 750 Republican Street, Box 358058, Seattle WA 98109; Phone: (206) 221-0594;; or Jianhai Du, One Medical Center Dr, PO Box 9193, West Virginia University Eye Institute, Morgantown, WV 26505; Phone: (304)-598-6903; Fax: (304)-598-6928;.

**This PDF file includes:**

Figure S1-S5

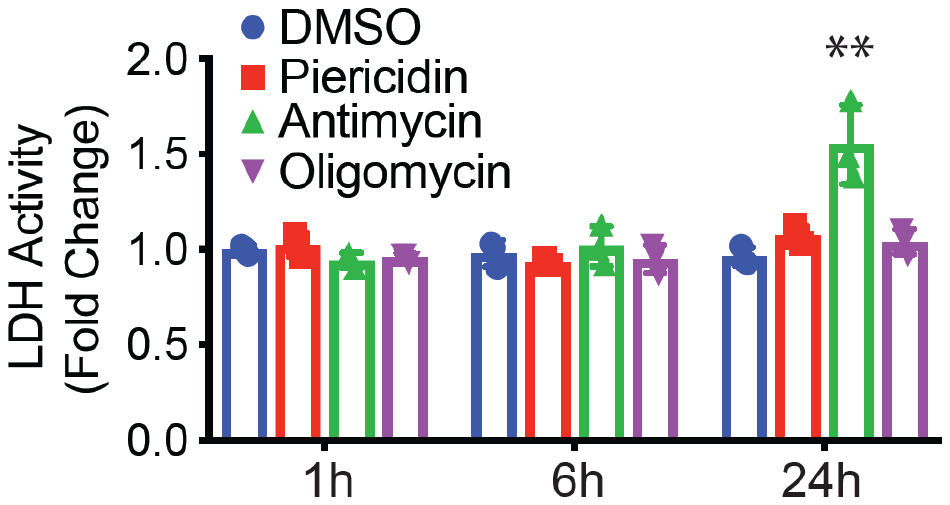

**Figure S1. The impact of mitochondrial inhibitors on LDH activity.** The data were presented as fold changes of Δ absorbance density at 340nm over the groups with DMSO at different time points. N=3. **P<0.01 vs. the groups treated with DMSO.

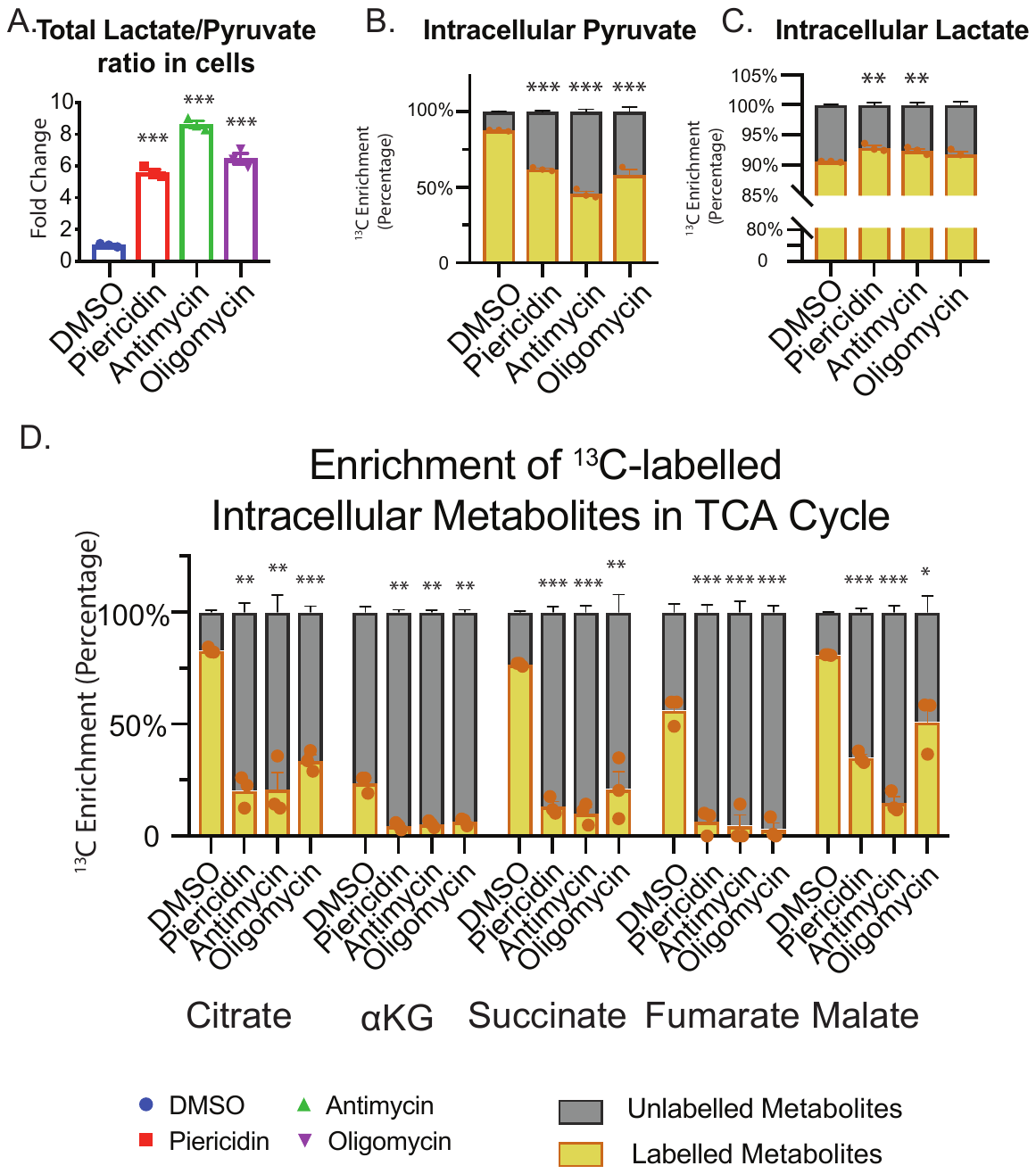

**Figure S2. Inhibition of mitochondrial respiration impairs the generation of glycolytic metabolites and TCA cycle intermediates from ^13^C-labeled glucose**. **(A)** Lactate/pyruvate ratio over the groups with DMSO. **(B-D)** Enrichment of ^13^C-labeled intracellular metabolites in glycolysis and the TCA cycle. N=3. *P<0.05, **P<0.01, *P<0.001 vs. the groups treated with DMSO.

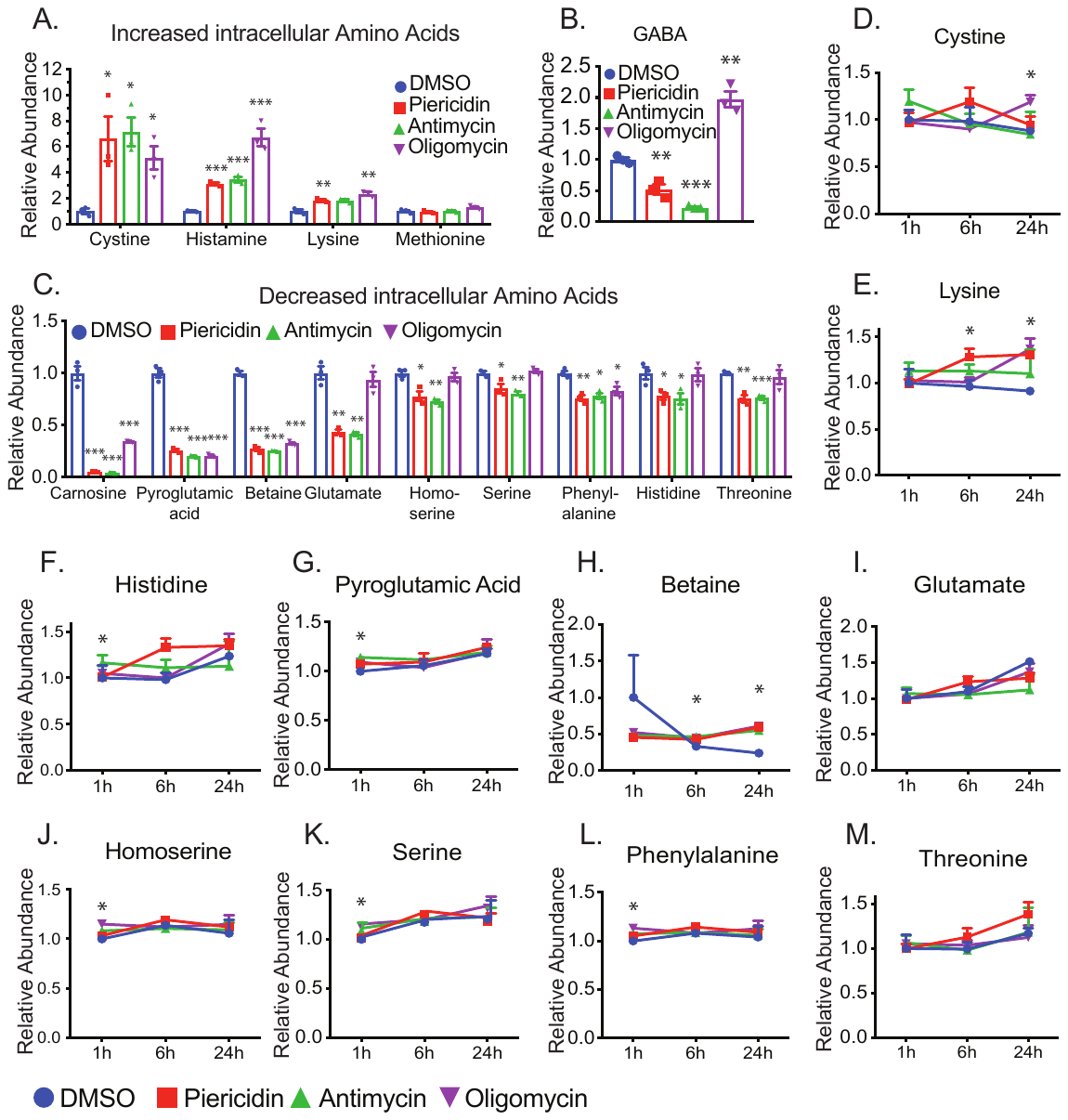

**Figure S3. Mitochondrial inhibition impairs the metabolism of amino acids.** **(A-C)** The relative abundance of intracellular amino acids. **(D-M)** The relative abundance of amino acids in media at different time points. N=3. *P<0.05, **P<0.01, *P<0.001 vs. the groups treated with DMSO.

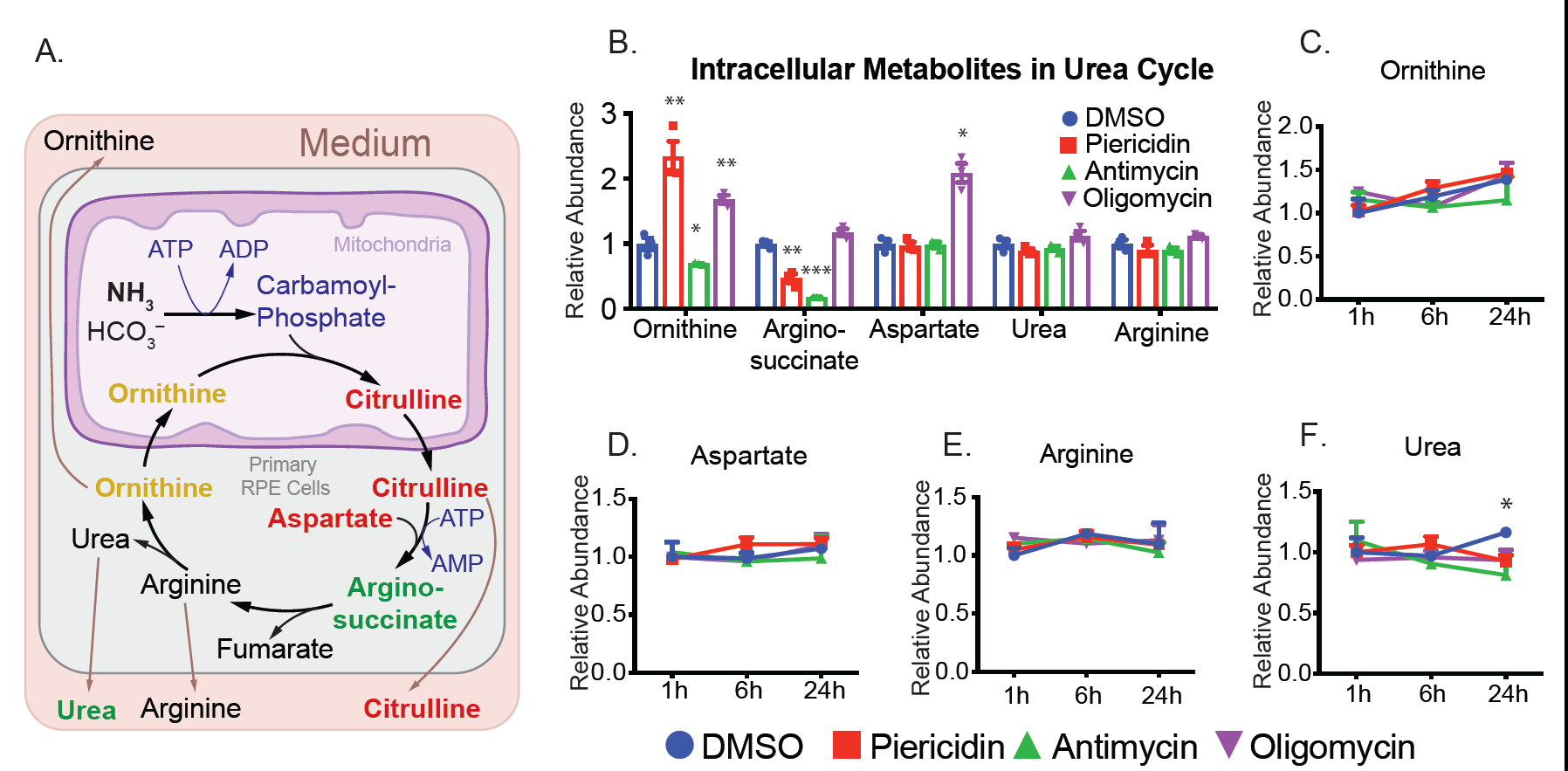

**Figure S4. Mitochondrial inhibition impairs the urea cycle.** **(A)** Schematic for the urea cycle. The color of metabolites represents the changes in relative abundance with mitochondrial inhibitors: red for the increased metabolites, green for decreased metabolites, orange for metabolites with mixed changes, and black for no change or not detected.) **(B)** Relative abundance of intracellular metabolites in the urea cycle. **(C-F)** The relative abundance of metabolites in the urea cycle in media at different time points. N=3. *P<0.05, **P<0.01, *P<0.001 vs the groups treated with DMSO.

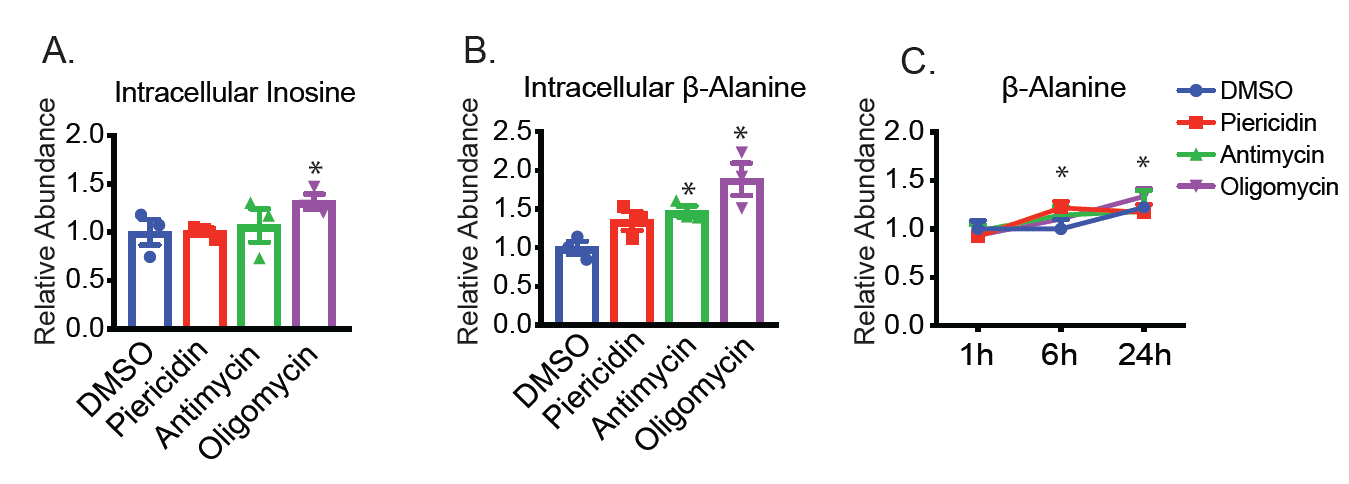

**Figure S5. The impact of mitochondrial inhibition on the metabolism of nucleotides.** **(A-B)** The relative abundance of intracellular inosine and β-alanine. **(C)** The relative abundance of β-alanine in media at different time points. N=3. *P<0.05 vs. the groups treated with DMSO.
